## Supplementary Information for "Sedimentation rate and organic matter dynamics shape microbiomes across a continental margin"

**Running Title:**	Sediment microbiomes across a continental margin

**Contents**

**Supplementary Tables**

**Tables S1-S3.** Sediment-depths explored along SSK42/5, SSK42/6 and SSK42/9, respectively, for microbiological analyses, including those based on duplicate metagenome sequencing (run accession numbers given).

**Tables S4-S8.** Compositions of the genomic sequence databases curated for sulfate-reducing bacteria and archaea, methanogenic archaea, anaerobic methanotrophs, acetogenic bacteria and anaerobic sulfur-oxidizing bacteria, respectively, for use in Bowtie2-based mapping of metatranscriptomic reads.

**Table S9.** Spearman correlation coefficients (*ρ*) and corresponding probability (*P*) values determined between the relative abundances of individual phyla/classes and sediment-depth, in the three cores SSK42/5, SSK42/6 and SSK42/9.

**Table S10.** Calculation of diversity indices for the individual sediment-samples of SSK42/5 using the phylum-level classification of their metagenomic data. Since this Table is more than one page long it has been provided as an Excel sheet named Table S10, within the Excel Workbook named Supplementary Dataset.

**Table S11.** Calculation of diversity indices for the individual sediment-samples of SSK42/6 using the phylum-level classification of their metagenomic data. Since this Table is more than one page long it has been provided as an Excel sheet named Table S11, within the Excel Workbook named Supplementary Dataset.

**Table S12.** Calculation of diversity indices for the individual sediment-samples of SSK42/9 using the phylum-level classification of their metagenomic data. Since this Table is more than one page long it has been provided as an Excel sheet named Table S12, within the Excel Workbook named Supplementary Dataset.

**Table S13.** Spearman correlation coefficients (*ρ*) and corresponding probability (*P*) values determined between the individual diversity indices and sediment-depth, in the three cores SSK42/5, SSK42/6 and SSK42/9.

**Table S14.** Number of homologs identified for the various structural genes associated with sulfate reduction, within the metagenome assemblies obtained for the individual sediment-samples of SSK42/5 and SSK42/6. Since this Table is more than one page long it has been provided as an Excel sheet named Table S14, within the Excel Workbook named Supplementary Dataset.

**Table S15.** Number of homologs identified for the various structural genes associated with methanogenesis, within the metagenome assemblies obtained for the individual sediment-samples of SSK42/5 and SSK42/6. Since this Table is more than one page long it has been provided as an Excel sheet named Table S15, within the Excel Workbook named Supplementary Dataset.

**Table S16.** Number of homologs identified for the various structural genes associated with acetogenesis, within the metagenome assemblies obtained for the individual sediment-samples of SSK42/5 and SSK42/6. Since this Table is more than one page long it has been provided as an Excel sheet named Table S16, within the Excel Workbook named Supplementary Dataset.

**Table S17.** Number of homologs identified for the various structural genes associated with anaerobic sulfur oxidation, within the metagenome assemblies obtained for the individual sediment-samples of SSK42/5 and SSK42/6. Since this Table is more than one page long it has been provided as an Excel sheet named Table S17, within the Excel Workbook named Supplementary Dataset.

**Table S18.** Number of homologs identified for the various structural genes associated with sulfate reduction, within the metagenome assemblies obtained for the individual sediment-samples of SSK42/9. Since this Table is more than one page long it has been provided as an Excel sheet named Table S18, within the Excel Workbook named Supplementary Dataset.

**Table S19.** Number of homologs identified for the various structural genes associated with methanogenesis, within the metagenome assemblies obtained for the individual sediment-samples of SSK42/9. Since this Table is more than one page long it has been provided as an Excel sheet named Table S19, within the Excel Workbook named Supplementary Dataset.

**Table S20.** Number of homologs identified for the various structural genes associated with acetogenesis, within the metagenome assemblies obtained for the individual sediment-samples of SSK42/9. Since this Table is more than one page long it has been provided as an Excel sheet named Table S20, within the Excel Workbook named Supplementary Dataset.

**Table S21.** Number of homologs identified for the various structural genes associated with anaerobic sulfur oxidation, within the metagenome assemblies obtained for the individual sediment-samples of SSK42/9. Since this Table is more than one page long it has been provided as an Excel sheet named Table S21, within the Excel Workbook named Supplementary Dataset.

**Table S22.** Spearman correlation coefficients (*ρ*) and corresponding probability (*P*) values determined between the relative abundances of individual metabolic-types and sediment-depth, in the three cores SSK42/5, SSK42/6 and SSK42/9.

**Table S23.** Spearman correlation coefficients (*ρ*) and corresponding probability (*P*) values determined pair-wise between the relative abundances of different metabolic-types, in the three cores SSK42/5, SSK42/6 and SSK42/9.

**Table S24.** Spearman correlation coefficients (*ρ*) and corresponding probability (*P*) values calculated pair-wise between the relative abundances of individual sulfur-metabolizing types, concentrations of different sulfur species and sediment-depth, in the three cores SSK42/5, SSK42/6 and SSK42/9.

**Table S25.** Number of homologs identified for the various structural genes associated with sulfate reduction, within the contigs assembled from the metatranscriptomic sequence data obtained for 0 cmbsf of SSK42/5 and 2 cmbsf of SSK42/6. Since this Table is more than one page long it has been provided as an Excel sheet named Table S25, within the Excel Workbook named Supplementary Dataset.

**Table S26.** Number of homologs identified for the various structural genes associated with methanogenesis, within the contigs assembled from the metatranscriptomic sequence data obtained for 0 cmbsf of SSK42/5 and 2 cmbsf of SSK42/6. Since this Table is more than one page long it has been provided as an Excel sheet named Table S26, within the Excel Workbook named Supplementary Dataset.

**Table S27.** Number of homologs identified for the various structural genes associated with acetogenesis, within the contigs assembled from the metatranscriptomic sequence data obtained for 0 cmbsf of SSK42/5 and 2 cmbsf of SSK42/6. Since this Table is more than one page long it has been provided as an Excel sheet named Table S27, within the Excel Workbook named Supplementary Dataset.

**Table S28.** Number of homologs identified for the various structural genes associated with anaerobic sulfur oxidation, within the contigs assembled from the metatranscriptomic sequence data obtained for 0 cmbsf of SSK42/5 and 2 cmbsf of SSK42/6. Since this Table is more than one page long it has been provided as an Excel sheet named Table S28, within the Excel Workbook named Supplementary Dataset.

**Tables S29 and S30.** Sulfate-reducing genera that were found to be most abundant along SSK42/5 and SSK42/6, respectively, according to the taxonomic classification of metagenomic reads (only those entities which had ≥ 5% relative abundance among all sulfate-reducing genera identified in a given sediment-sample have been included).

**Tables S31 and S32.** Methanogenic genera that were found to be most abundant along SSK42/5 and SSK42/6, respectively, according to the taxonomic classification of metagenomic reads (only those entities which had ≥ 5% relative abundance among all methane-producing genera identified in a given sediment-sample have been included).

**References used in the Supplementary Tables**

**Supplementary Notes**

**Supplementary Note 1.** Referred citations for the sulfate-reducing bacterial and archaeal taxa, summing up the mean relative abundances of which gave the measure of prevalence of sulfate-reducers within a sediment community of SSK42/5, SSK42/6 or SSK42/9.

**Supplementary Note 2.** Referred citations for the anaerobically sulfur-oxidizing chemolithotrophic bacteria (ANSOB), summing up the mean relative abundances of which gave the measure of prevalence of ANSOB within a sediment community of SSK42/5, SSK42/6 or SSK42/9.

**Supplementary Tables**

**Table S1.** Sediment-depths explored along SSK42/5 for microbiological analyses, including those based on duplicate metagenome sequencing (run accession numbers given).

| **Sediment-depths explored**  **(in cmbsf)** | **BioSample accession number** | **Sample-replicate** | **Run accession number** |
| --- | --- | --- | --- |
| 0 | SAMN04442175 | 1st replicate | SRR3646127 |
|  |  | 2nd replicate | SRR3646128 |
| 15 | SAMN04442176 | 1st replicate | SRR3646129 |
|  |  | 2nd replicate | SRR3646130 |
| 45 | SAMN04442177 | 1st replicate | SRR3646131 |
|  |  | 2nd replicate | SRR3646132 |
| 60 | SAMN04442178 | 1st replicate | SRR3646144 |
|  |  | 2nd replicate | SRR3646145 |
| 90 | SAMN04442179 | 1st replicate | SRR3646147 |
|  |  | 2nd replicate | SRR3646148 |
| 120 | SAMN04442180 | 1st replicate | SRR3646150 |
|  |  | 2nd replicate | SRR3646151 |
| 140 | SAMN04442181 | 1st replicate | SRR3646152 |
|  |  | 2nd replicate | SRR3646153 |
| 160 | SAMN04442182 | 1st replicate | SRR3646155 |
|  |  | 2nd replicate | SRR3646156 |
| 190 | SAMN04442183 | 1st replicate | SRR3646157 |
|  |  | 2nd replicate | SRR3646158 |
| 220 | SAMN04442184 | 1st replicate | SRR3646160 |
|  |  | 2nd replicate | SRR3646161 |
| 260 | SAMN04442185 | 1st replicate | SRR3646162 |
|  |  | 2nd replicate | SRR3646163 |
| 295 | SAMN04442186 | 1st replicate | SRR3646164 |
|  |  | 2nd replicate | SRR3646165 |

**Table S2.** Sediment-depths explored along SSK42/6 for microbiological analyses, including those based on duplicate metagenome sequencing (run accession numbers given).

| **Sediment-depths explored**  **(in cmbsf)** | **BioSample accession number** | **Sample-fraction** | **Run accession number** |
| --- | --- | --- | --- |
| 2 | SAMN04442187 | 1st replicate | SRR3570036 |
|  |  | 2nd replicate | SRR3570038 |
| 30 | SAMN04442189 | 1st replicate | SRR3577067 |
|  |  | 2nd replicate | SRR3577068 |
| 45 | SAMN04442190 | 1st replicate | SRR3577070 |
|  |  | 2nd replicate | SRR3577071 |
| 60 | SAMN04442191 | 1st replicate | SRR3577073 |
|  |  | 2nd replicate | SRR3577076 |
| 75 | SAMN04442192 | 1st replicate | SRR3577078 |
|  |  | 2nd replicate | SRR3577079 |
| 90 | SAMN04442193 | 1st replicate | SRR3577081 |
|  |  | 2nd replicate | SRR3577082 |
| 120 | SAMN04442195 | 1st replicate | SRR3577086 |
|  |  | 2nd replicate | SRR3577087 |
| 135 | SAMN04442196 | 1st replicate | SRR3577090 |
|  |  | 2nd replicate | SRR3577311 |
| 175 | SAMN04442199 | 1st replicate | SRR3577337 |
|  |  | 2nd replicate | SRR3577338 |
| 220 | SAMN04442202 | 1st replicate | SRR3577341 |
|  |  | 2nd replicate | SRR3577343 |
| 250 | SAMN04442204 | 1st replicate | SRR3577344 |
|  |  | 2nd replicate | SRR3577345 |
| 265 | SAMN04442205 | 1st replicate | SRR3577347 |
|  |  | 2nd replicate | SRR3577349 |
| 275 | SAMN04442207 | 1st replicate | SRR3577350 |
|  |  | 2nd replicate | SRR3577351 |

**Table S3.** Sediment-depths explored along SSK42/9 for microbiological analyses, including those based on duplicate metagenome sequencing (run accession numbers given).

| **Sediment-depths explored**  **(in cmbsf)** | **BioSample accession number** | **Sample-fraction** | **Run accession number** |
| --- | --- | --- | --- |
| 0 | SAMN04442233 | 1st replicate | SRR3872933 |
|  |  | 2nd replicate | SRR3872934 |
| 19 | SAMN04442236 | 1st replicate | SRR3884351 |
|  |  | 2nd replicate | SRR3884355 |
| 50 | SAMN04442238 | 1st replicate | SRR3884357 |
|  |  | 2nd replicate | SRR3884359 |
| 115 | SAMN04442241 | 1st replicate | SRR3884468 |
|  |  | 2nd replicate | SRR3884472 |
| 125 | SAMN04442242 | 1st replicate | SRR3884479 |
|  |  | 2nd replicate | SRR3884488 |
| 145 | SAMN04442244 | 1st replicate | SRR3884538 |
|  |  | 2nd replicate | SRR3884540 |
| 155 | SAMN04442245 | 1st replicate | SRR3884542 |
|  |  | 2nd replicate | SRR3884544 |
| 180 | SAMN04442246 | 1st replicate | SRR3884546 |
|  |  | 2nd replicate | SRR3884547 |
| 225 | SAMN04442248 | 1st replicate | SRR3884548 |
|  |  | 2nd replicate | SRR3884552 |
| 255 | SAMN04442250 | 1st replicate | SRR3884553 |
|  |  | 2nd replicate | SRR3884554 |

**Table S4.** Composition of the genomic sequence database curated for sulfate-reducing bacteria and archaea, and used in Bowtie2-based mapping of metatranscriptomic reads.

| **Serial number** | **Genera^1^ included in the curated database** | **Strain whose genome sequence was included in the database** | **GenBank accession number of the genome sequence included in the database** | **Genome size**  **(in bp)** | **Number of CDS present in the genome** |
| --- | --- | --- | --- | --- | --- |
| 1. | Archaeoglobus fulgidus | DSM 4304 | NC_000917, NZ_AE000943-NZ_AE001114 | 2,178,400 | 2,369 |
| 2. | *Desulfacinum infernum* | DSM 9756 | NZ_FQVB00000000 | 4,239,813 | 3,584 |
| 3. | *Desulfarculus baarsii* | DSM 2075 | NC_014365 | 3,655,731 | 3,244 |
| 4. | *Desulfatibacillum aliphaticivorans* | DSM 15576 | NZ_AUCT01000000 | 6,466,399 | 5,264 |
| 5. | *Desulfatiglans anilini* | DSM 4660 | NZ_AULM01000000 | 4,664,663 | 3,943 |
| 6. | *Desulfatirhabdium butyrativorans* | DSM 18734 | NZ_AUCU01000000 | 4,475,774 | 3,852 |
| 7. | *Desulfatitalea tepidiphila* | S28bF | NZ_BCAG01000000 | 5,614,477 | 4,858 |
| 8. | *Desulfitibacter alkalitolerans* | DSM 16504 | NZ_JHVU01000000 | 4,341,391 | 4,096 |
| 9. | *Desulfitobacterium dehalogenans* | ATCC 51507 | NC_018017,  NZ_AGJH01000000,NZ_AGJH01000001-NZ_AGJH01000066 | 4,321,753 | 3,974 |
| 10. | *Desulfobacca acetoxidans* | DSM 11109 | NC_015388 | 3,282,536 | 2,888 |
| 11. | *Desulfobacter postgatei* | 2ac9 | NZ_AGJR02000000 | 3,972,458 | 3,485 |
| 12. | *Desulfobacterium autotrophicum*^2^ | HRM2 | NC_012108 | 5,589,073 | 4,835 |
| 13. | *Desulfobacterium vacuolatum*^2^ | DSM 3385 | NZ_FWXY00000000 | 5,035,353 | 4,050 |
| 14. | *Desulfobacula toluolica* | Tol2 | NC_018645 | 5,197,905 | 4,545 |
| 15. | *Desulfobulbus propionicus* | DSM 2032 | CP002364 | 3,851,869 | 3,283 |
| 16. | *Desulfocapsa sulfexigens*^3^ | DSM 10523 | NC_020304 | 3,986,761 | 3,522 |
| 17. | *Desulfocarbo indianensis* | SCBM | NZ_LBMP00000000 | 5,114,003 | 4,383 |
| 18. | *Desulfococcus multivorans* | DSM 2059 | NZ_CP015381 | 4,455,399 | 3,813 |
| 19. | *Desulfocurvus vexinensis* | DSM 17965 | NZ_JAEX00000000 | 3,623,604 | 3,200 |
| 20. | *Desulfofaba hansenii*^4^ | P1 | NZ_PESK00000000 | 6,710,842 | 5,239 |
| 21. | *Desulfofustis glycolicus* | DS 9705 | NZ_FQXS00000000 | 4,975,017 | 4,297 |
| 22. | *Desulfohalobium retbaense* | DSM 5692 | NC_013223, NZ_ABTO01000000,NZ_ABTO01000001-NZ_ABTO01000032 | 2,864,304 | 2,488 |
| 23. | *Desulfoluna spongiiphila*^5^ | AA1 | NZ_FMUX00000000 | 6,530,021 | 5,203 |
| 24. | *Desulfomicrobium baculatum* | DSM 4028 | NC_013173,  NZ_ABTP01000000,NZ_ABTP01000001- NZ_ABTP01000013 | 3,942,657 | 3,395 |
| 25. | *Desulfomonile tiedjei* | DSM 6799 | NC_018025 | 6,500,104 | 5,551 |
| 26. | *Desulfonatronospira thiodismutans* | ASO3-1 | NZ_ACJN00000000 | 4,107,394 | 3,729 |
| 27. | *Desulfonatronovibrio hydrogenovorans* | DSM 9292 | NZ_JMKT00000000 | 2,939,095 | 2,761 |
| 28. | *Desulfonatronum lacustre* | DSM 10312 | NZ_JAFE00000000 | 3,761,151 | 3,303 |
| 29. | *Desulfonauticus submarines* | DSM 15269 | NZ_FNIN00000000 | 2,100,696 | 2,014 |
| 30. | *Desulfonispora thiosulfatigenes* | DSM 11270 | NZ_FWWT00000000 | 2,412,168 | 2,345 |
| 31. | *Desulfopila aestuarii* | DSM 18488 | NZ_FRFE00000000 | 6,065,581 | 5,073 |
| 32. | *Desulfoplanes formicivorans* | Pf12B | NZ_BDFE00000000 | 3,000,887 | 2,628 |
| 33. | *Desulforegula conservatrix* | Mb1Pa | NZ_AUEY00000000 | 4,464,516 | 3,803 |
| 34. | *Desulforhopalus singaporensis*^6^ | DSM 12130 | NZ_FNJI00000000 | 5,006,845 | 4,203 |
| 35. | *Desulfosarcina cetonica*^7^ | JCM 12296 | NZ_BBCC00000000 | 7,092,334 | 5,582 |
| 36. | *Desulfospira joergensenii* | DSM 10085 | NZ_ATUG00000000 | 6,119,470 | 5,490 |
| 37. | *Desulfosporosinus orientis* | DSM 765 | NC_016584 | 5,863,081 | 5,229 |
| 38. | *Desulfotalea psychrophila* | LSv54 | NC_006138 | 3,523,383 | 3,128 |
| 39. | *Desulfothermus okinawensis*^8^ | JCM 13304 | BBBM00000000 | 2,774,183 | ND |
| 40. | *Desulfotignum balticum* | DSM 7044 | NZ_ATWO00000000 | 5,118,755 | 4,623 |
| 41. | *Desulfotomaculum nigrificans* | CO-1-SRB | NC_015565 | 2,892,255 | 2,647 |
| 42. | *Desulfovermiculus halophilus* | DSM 18834 | NZ_JIAK00000000 | 3,241,624 | 2,903 |
| 43. | *Desulfovibrio desulfuricans subsp. Desulfuricans*^9^ | ATCC 27774 | NC_011883,  NZ_ACCT01000000,NZ_ACCT01000001-NZ_ACCT01000010 | 2,873,437 | 2,406 |
| 44. | *Desulfovibrio desulfuricans*^9^ | ND132 | NC_016803, NZ_AEUJ01000000,NZ_AEUJ01000001- NZ_AEUJ01000007, NZ_CM001077 | 3,858,580 | 3,423 |
| 45. | *Desulfovirgula thermocuniculi* | DSM 16036 | NZ_AUBR00000000 | 3,054,371 | 3,031 |
| 46. | *Desulfurella acetivorans* | A63 | NZ_CP007051 | 1,816,487 | 1,814 |
| 47. | *Desulfurispirillum indicum*^10^ | S5 | NC_014836,  NZ_ADGU01000000,NZ_ADGU01000001-,NZ_ADGU01000113 | 2,928,377 | 2,576 |
| 48. | *Desulfurispora thermophila* | DSM 16022 | NZ_AQWN00000000 | 2,821,579 | 2,661 |
| 49. | *Desulfurivibrio alkaliphilus* | AHT2 | NC_014216, NZ_ACYL01000000, NZ_ACYL01000001,-NZ_ACYL01000066 | 3,097,763 | 2,643 |
| 50. | *Desulfurobacterium thermolithotrophum* | DSM 11699 | NC_015185 | 1,541,968 | 1,508 |
| 51. | *Desulfurococcus mucosus* | DSM 2162 | NC_014961 | 1,314,639 | 1,345 |
| 52. | *Desulfuromonas acetoxidans* | DSM 684 | NZ_AAEW00000000 | 3,828,328 | 3,241 |
| 53. | *Desulfuromusa kysingii* | DSM 7343 | NZ_FNQN00000000 | 3,742,394 | 3,300 |
| 54. | *Dethiobacter alkaliphilus* | AHT 1 | NZ_ACJM00000000 | 3,116,746 | 3,067 |
| 55. | *Dethiosulfatibacter aminovorans* | DSM 17477 | NZ_FQZL00000000 | 4,031,536 | 3,531 |
| 56. | *Dethiosulfovibrio peptidovorans* | DSM 11002 | NZ_ABTR00000000 | 2,576,359 | 2,399 |
| 57. | *Thermodesulfatator indicus* | DSM 15286 | NC_015681 | 2,322,224 | 2,226 |
| 58. | *Thermodesulfobacterium commune* | DSM 2178 | NZ_CP008796 | 1,764,045 | 1,702 |
| 59. | *Thermodesulfobium narugense* | DSM 14796 | NC_015499 | 1,898,865 | 1,825 |
| 60. | *Thermodesulforhabdus norvegica* | DSM 9990 | NZ_FOUU00000000 | 2,801,945 | 2,631 |

^1^ All these genera have sulfur reduction reported from its members.

^2^ *Desulfobacterium* contain 9 species; the type species is *Desulfobacterium indolicum* but its genome sequence is not available so all the genome sequences available for different species of *Desulfobacterium* was included in this database.

^3^ *Desulfocapsa* contains two species: the type species is *Desulfocapsa* *thiozymogenes*; but its genome sequence is not available; so the sequence of *D*. sulfexigens DSM10523was included in this database.

^4^ *Desulfofaba* contains three species: the type species is *Desulfofaba gelida*; but its genome sequence is not available. The only available genome sequence from this genera is the genome sequence of *D. hansenii* P1, so it is included in this database.

^5^ *Desulfoluna* contains two species: the type species is *Desulfoluna butyratoxydans*; but its genome sequence is not available; so the sequence of *D*. *spongiiphila* AA1 was included in this database.

^6^ *Desulforhopalus* contains two species: the type species is *Desulforhopalus vacuolatus*; but its genome sequence is not available; so the sequence of *D*. singaporensis DSM12130 was included in this database.

^7^ *Desulfosarcina* contains five species: the type species is *Desulfosarcina variabilis*; but its genome sequence is not available. The only available genome sequence from this genera is the genome sequence of *D. cetonica* JCM 12296, so it is included in this database.

^8^ *Desulfothermus* contains two species: the type species is *Desulfothermus naphthae*; but its genome sequence is not available; so the sequence of *D*. *okinawensis* JCM 13304 was included in this database.

^9^ The type species of *Desulfovibrio* is *Desulfovibrio desulfuricans*. Genome sequence is available for two strains of *D. desulfuricans*, both the sequence was included in this database.

^10^ *Desulfurispirillum* contains two species: the type species is *Desulfurispirillum alkaliphilum*; but its genome sequence is not available; so the sequence of *D*. *indicum* S5 was included in this database.

**Table S5.** Composition of the genomic sequence database curated for methanogenic archaea, and used in Bowtie2-based mapping of metatranscriptomic reads.

| **Serial number** | **Genera^1^ included in the curated database** | **Strain whose genome sequence was included in the database** | **GenBank accession number of the genome sequence included in the database** | **Genome size**  **(in bp)** | **Number of CDS present in the genome** |
| --- | --- | --- | --- | --- | --- |
| 1. | *Methanobacterium formicicum* | DSM 3637 | NZ_AMPO00000000 | 2,684,267 | 2,476 |
| 2. | *Methanobrevibacter ruminantium* | M1 | NC_013790 | 2,937,203 | 2,143 |
| 3. | *Methanocaldococcus jannaschii* | DSM 2661 | NC_000909  NZ_U67459-NZ_U67608 | 1,664,970 | 1,762 |
| 4. | *Methanocella paludicola* | SANAE | AP011532 | 2,957,635 | 3,004 |
| 5. | *Methanococcoides methylutens*^2^ | DSM 2657 | NZ_JRHO00000000 | 2,508,511 | 2,355 |
| 6. | *Methanococcoides methylutens*^2^ | MM1 | NZ_CP009518 | 2,394,636 | 2,245 |
| 7. | *Methanocorpusculum parvum* | XII | NZ_LMVO00000000 | 1,709,133 | 1,688 |
| 8. | *Methanoculleus bourgensis* | MS2T | NC_018227 | 2,789,774 | 2,538 |
| 9. | *Methanofollis ethanolicus*^3^ | HASU | NZ_BCNW00000000 | 2,713,913 | 2,554 |
| 10. | *Methanofollis liminatans*^3^ | DSM 4140 | NZ_CM001555 NZ_AGCO01000000 | 2,475,100 | 2,396 |
| 11. | *Methanogenium cariaci* | JCM 10550 | NZ_BBBG00000000 | 2,686,643 | 2,258 |
| 12. | *Methanohalobium evestigatum* | Z-7303 | NC_014253 | 2,242,317 | 2,267 |
| 13. | *Methanohalophilus mahii* | DSM 5219 | NC_014002 | 2,012,424 | 1,955 |
| 14. | *Methanolacinia paynteri* | DSM 2545 | NZ_AXDV00000000 | 2,785,552 | 2,664 |
| 15. | *Methanolinea tarda* | NOBI-1 | NZ_AGIY02000001 NZ_AGIY00000000 | 2,052,856 | 2,010 |
| 16. | *Methanolobus tindarius* | DSM 2278 | NZ_AZAJ00000000 | 3,151,883 | 2,886 |
| 17. | *Methanomassiliicoccus luminyensis* | B10 | NZ_CAJE00000000 | 2,620,233 | 2,555 |
| 18. | *Methanomethylovorans hollandica* | DSM 15978 | NC_019977 | 2,428,904 | 2,525 |
| 19. | *Methanomicrobium mobile* | BP | NZ_JOMF00000000 | 1,711,791 | 1,617 |
| 20. | *Methanoplanus limicola* | DSM 2279 | NZ_CM001436 NZ_AHKP01000000 | 3,200,946 | 2,967 |
| 21. | *Methanopyrus kandleri* | AV19 | AE009439 AE010302-AE010458 | 1,694,969 | 1,687 |
| 22. | *Methanoregula boonei* | 6A8 | NC_009712 | 2,542,943 | 2,513 |
| 23. | *Methanosalsum zhilinae* | DSM 4017 | NC_015676 | 2,138,444 | 1,955 |
| 24. | *Methanosarcina acetivorans*^4^ | C2A | NC_003552 NZ_AE010656-NZ_AE011189 | 5,751,492 | 4,567 |
| 25. | *Methanosarcina barkeri*^4^ | MS | NZ_CP009528 | 4,533,209 | 3,481 |
| 26. | *Methanosarcina barkeri*^4^ | 3 | NZ_CP009517 | 4,560,446 | 3,589 |
| 27. | *Methanosarcina flavescens*^4^ | E03.2 | NZ_LKAZ00000000 | 3,268,085 | 2,652 |
| 28. | *Methanosarcina horonobensis*^4^ | HB-1 | NZ_CP009516 | 5,018,607 | 4,144 |
| 29. | *Methanosarcina lacustris*^4^ | Z-7289 | NZ_CP009515 | 4,139,808 | 3,264 |
| 30. | *Methanosarcina mazei*^4^ | S-6 | NZ_CP009512 | 4,142,816 | 3,359 |
| 31. | *Methanosarcina siciliae*^4^ | T4/M | NZ_CP009506 | 5,017,558 | 3,930 |
| 32. | *Methanosarcina soligelidi*^4^ | SMA-21 | NZ_JQLR00000000 | 4,064,496 | 3,257 |
| 33. | *Methanosarcina spelaei*^4^ | MC-15 | NZ_LMVP00000000 | 5,088,600 | 3,774 |
| 34. | *Methanosarcina thermophila*^4^ | TM-1 | NZ_CP009501 | 3,127,379 | 2,597 |
| 35. | *Methanosarcina vacuolata*^4^ | Z-761 | NZ_CP009520 | 4,505,752 | 3,538 |
| 36. | *Methanosphaera stadtmanae* | DSM 3091 | NC_007681 | 1,767,403 | 1,518 |
| 37. | *Methanosphaerula palustris* | E1-9c | NC_011832  NZ_ABZB01000000 NZ_ABZB01000001 NZ_ABZB01000002  NZ_ABZB01000003 NZ_ABZB01000004 NZ_ABZB01000005 NZ_ABZB01000006  NZ_ABZB01000007 | 2,922,917 | 2,696 |
| 38. | *Methanospirillum hungatei* | JF-1 | NC_007796 NZ_AALU01000000 NZ_AALU01000001-NZ_AALU01000090 | 3,544,738 | 3,294 |
| 39. | *Methanothermobacter thermautotrophicus* | Delta H | NC_000916 NZ_AE000795-NZ_AE000942 | 1,751,377 | 1,756 |
| 40. | *Methanothermococcus thermolithotrophicus* | DSM 2095 | NZ_AQXV00000000 | 1,686,930 | 1,624 |
| 41. | *Methanothermus fervidus* | DSM 2088 | NC_014658 | 1,243,342 | 1,296 |
| 42. | *Methanosaeta concilii* | GP-6 | NC_015416 | 3,008,626 | 2,899 |
| 43. | *Methanotorris igneus* | Kol 5 | NC_015562 | 1,854,197 | 1,751 |
| 44. | *Methermicoccus shengliensis* | DSM 18856 | NZ_JONQ00000000 | 1,511,083 | 1,558 |

^1^ All these genera have methanogenesis reported from its members.

^2^ Type strain of *Methanococcoides methylutens* is DSM 2657. Genome sequence of *Methanococcoides methylutens* MM1 was also available. So both the genome sequences were included.

^3^ *Methanofollis* contain 5 species; the type species is *Methanofollis tationis* but its genome sequence is not available so all the genome sequences available for different species of *Methanofollis* was included in this database.

^4^ *Methanosarcina* contain 16 species; the type species is *Methanosarcina methanica* but its genome sequence is not available so all the genome sequences available for different species of *Methanosarcina* was included in this database.

**Table S6.** Composition of the genomic sequence database curated for anaerobic methanotrophs, and used in Bowtie2-based mapping of metatranscriptomic reads.

| **Serial number** | **Genera^1^ included in the curated database** | **Strain whose genome sequence was included in the database** | **GenBank accession number of the genome sequence included in the database** | **Genome size**  **(in bp)** | **Number of CDS present in the genome** |
| --- | --- | --- | --- | --- | --- |
| 1. | Uncultured archaeon | clone GZfos10C7 | AY714814 | 36,996 | ND |
| 2. | Uncultured archaeon | clone GZfos11H11 | AY714816 | 36,155 | ND |
| 3. | Uncultured archaeon | clone GZfos12E1 | AY714817 | 35,297 | ND |
| 4. | Uncultured archaeon | clone GZfos13E1 | AY714819 | 35,492 | ND |
| 5. | Uncultured archaeon | clone GZfos17A3 | AY714821 | 37,604 | ND |
| 6. | Uncultured archaeon | clone GZfos18B6 | AY714825 | 37,726 | ND |
| 7. | Uncultured archaeon | clone GZfos18C8 | AY714826 | 39,495 | ND |
| 8. | Uncultured archaeon | clone GZfos19C7 | AY714830 | 33,603 | ND |
| 9. | Uncultured archaeon | clone GZfos1C11 | AY714832 | 42,079 | ND |
| 10. | Uncultured archaeon | clone GZfos24D9 | AY714837 | 29,650 | ND |
| 11. | Uncultured archaeon | clone GZfos25D1 | AY714838 | 31,125 | ND |
| 12. | Uncultured archaeon | clone GZfos26B2 | AY714839 | 25,119 | ND |
| 13. | Uncultured archaeon | clone GZfos26E7 | AY714841 | 32,690 | ND |
| 14. | Uncultured archaeon | clone GZfos27G5 | AY714848 | 40,522 | ND |
| 15. | Uncultured archaeon | clone GZfos30H9 | AY714852 | 36,779 | ND |
| 16. | Uncultured archaeon | clone GZfos33H6 | AY714858 | 34,700 | ND |
| 17. | Uncultured archaeon | clone GZfos34G5 | AY714860 | 34,480 | ND |
| 18. | Uncultured archaeon | clone GZfos35D7 | AY714865 | 25,161 | ND |
| 19. | Uncultured archaeon | clone GZfos9C4 | AY714870 | 38,084 | ND |
| 20. | Candidatus *Methanoperedens nitroreducens* | ANME-2d | NZ_JMIY00000000 | 3,197,389 | 3,254 |
| 21. | Candidatus *Methanoperedens nitroreducens* | Mnv1 | NZ_FZMP00000000 | 3,524,740 | 3,494 |

^1^ All these genera have anaerobic methanotrophy reported from its members.

ND = Not defined

**Table S7.** Composition Composition of the genomic sequence database curated for acetogenic bacteria, and used in Bowtie2-based mapping of metatranscriptomic reads.

| **Serial number** | **Genera^1^ included in the curated database** | **Strain whose genome sequence was included in the database** | **GenBank accession number of the genome sequence included in the database** | **Genome size**  **(in bp)** | **Number of CDS present in the genome** |
| --- | --- | --- | --- | --- | --- |
| 1. | *Acetitomaculum ruminis* | DSM 5522 | NZ_FOJY00000000 | 3,078,669 | 2,537 |
| 2. | *Acetoanaerobium noterae* | ATCC 35199 | NZ_FUYN00000000 | 2,811,216 | 2,663 |
| 3. | *Acetobacterium woodii* | DSM 1030 | NC_016894 | 4,044,777 | 3,527 |
| 4. | *Acetohalobium arabaticum* | DSM 5501 | NC_014378 | 2,469,596 | 2,274 |
| 5. | *Acetonema longum* | DSM 6540 | NZ_AFGF00000000 | 4,323,011 | 3,847 |
| 6. | *Moorella thermoacetica* | ATCC 39073 | NC_007644 | 2,628,784 | 2,460 |
| 7. | *Oxobacter pfennigii* | DSM 3222 | NZ_LKET00000000 | 4,510,552 | 4,146 |
| 8. | *Ruminococcus flavefaciens* | ATCC 19208 | NZ_JAEF00000000 | 3,581,418 | 2,983 |
| 9. | *Sporomusa sphaeroides* | DSM 2875 | NZ_LSLJ00000000 | 4,971,384 | 4,463 |

^1^ All these genera have acetogenesis reported from its members.

**Supplementary Table S8.** Composition of the genomic sequence database curated for anaerobic sulfur-oxidizing bacteria, and used in Bowtie2-based mapping of metatranscriptomic reads.

| **Serial number** | **Genera^1^ included in the curated database** | **Strain whose genome sequence was included in the database** | **GenBank accession number of the genome sequence included in the database** | **Genome size**  **(in bp)** | **Number of CDS present in the genome** |
| --- | --- | --- | --- | --- | --- |
| 1. | *Beggiatoa alba* | B18LD | NZ_AHMA00000000 | 4,263,346 | 3,446 |
| 2. | *Sulfuricurvum kujiense* | DSM16994 | NC_014762 | 2,574,824 | 2,772 |
| 3. | *Sulfurimonas autotrophica* | DSM16294 | NC_014506 | 2,153,198 | 2,140 |
| 4. | *Sulfurovum lithotrophicum* | ATCCBAA-797 | NZ_CP011308 | 2,217,891 | 2,144 |
| 5. | *Thiobacillus thioparus* | DSM505 | NZ_ARDU00000000 | 3,200,073 | 3,071 |

^1^ All these genera have anaerobic sulfur oxidation reported from its members.

**Table S9.** Spearman correlation coefficients (*ρ*) and corresponding probability (*P*) values determined between the relative abundances of individual phyla/classes and sediment-depth, in the three cores SSK42/5, SSK42/6 and SSK42/9.

|  | **SSK42/5** | | **SSK42/6** | | **SSK42/9** | |
| --- | --- | --- | --- | --- | --- | --- |
| **Phyla** | ***ρ*** | ***P* value** | ***ρ*** | ***P* value** | ***ρ*** | ***P* value** |
| ***Crenarchaeota*** | -0.7 | 0.017305023 | -0.5 | 0.0673418 | 0.7 | 0.0311411 |
| ***Euryarchaeota*** | -0.7 | 0.007353207 | -0.5 | 0.0888743 | 0.7 | 0.0275141 |
| ***Thaumarchaeota*** | -0.6 | 0.027954553 | -0.5 | 0.0673418 | 0.6 | 0.087768 |
| ***Acidobacteria*** | -0.9 | 0.000192153 | -0.7 | 0.013897 | -0.4 | 0.2956041 |
| ***Actinobacteria*** | -0.9 | 0 | -0.4 | 0.2166298 | 0.03 | 0.9457098 |
| ***Bacteroidetes*** | -0.6 | 0.0457531 | -0.8 | 0.0016243 | -0.6 | 0.0802151 |
| ***Chloroflexi*** | -0.9 | 0.000597084 | -0.5 | 0.0610936 | 0.9 | 0.0019771 |
| ***Cyanobacteria*** | -0.3 | 0.403976659 | -0.4 | 0.1350282 | 0.2 | 0.4915555 |
| ***Firmicutes*** | -0.6 | 0.040002049 | -0.6 | 0.0448907 | 0.7 | 0.0241946 |
| ***Planctomycetes*** | -0.9 | 0 | -0.7 | 0.0052901 | -0.4 | 0.2180285 |
| ***Alphaproteobacteria*** | -0.3 | 0.286690914 | -0.4 | 0.1887717 | -0.2 | 0.4915555 |
| ***Betaproteobacteria*** | 0.6 | 0.03239475 | 0.4 | 0.2166298 | -0.6 | 0.0957916 |
| ***Deltaproteobacteria*** | -0.6 | 0.066629001 | -0.5 | 0.0775693 | -0.09 | 0.811417 |
| ***Gammaproteobacteria*** | 0.9 | 0 | 0.7 | 0.0120134 | -0.7 | 0.0311411 |
| ***Zetaproteobacteria*** | 0.8 | 0.006596543 | -0.1 | 0.7369512 | -0.6 | 0.0664691 |
| ***Thermotogae*** | -0.8 | 0.004659881 | -0.6 | 0.0340307 | 0.7 | 0.0350915 |
| ***Verrucomicrobia*** | -0.7 | 0.018825126 | -0.4 | 0.1350282 | 0.4 | 0.2956041 |

**Table S13.** Spearman correlation coefficients (*ρ*) and corresponding probability (*P*) values determined between the individual diversity indices and sediment-depth, in the three cores SSK42/5, SSK42/6 and SSK42/9.

|  | **SSK42/5** | | **SSK42/6** | | **SSK42/9** | |
| --- | --- | --- | --- | --- | --- | --- |
|  | ***ρ*** | ***P* value** | ***ρ*** | ***P* value** | ***ρ*** | ***P* value** |
| **D** | 0.8 | 0.0009 | 0.8 | 0.0026 | 0.2 | 0.6 |
| **H** | -0.8 | 0.0017 | -0.9 | 0.0001 | -0.07 | 0.9 |
| **E_H_** | -0.8 | 0.0017 | -0.9 | 0.0001 | -0.07 | 0.9 |

**Table S22.** Spearman correlation coefficients (*ρ*) and corresponding probability (*P*) values determined between the relative abundances of individual metabolic-types and sediment-depth, in the three cores SSK42/5, SSK42/6 and SSK42/9.

|  | **SSK42/5** | | **SSK42/6** | | **SSK42/9** | |
| --- | --- | --- | --- | --- | --- | --- |
|  | ***ρ*** | ***P* value** | ***ρ*** | ***P* value** | ***ρ*** | ***P* value** |
| **Sulfate-reducers** | -0.6 | 0.04 | -0.5 | 0.05 | 0.5 | 0.2 |
| **Methanogens** | -0.7 | 0.007 | -0.6 | 0.05 | 0.7 | 0.03 |
| **ANME** | -0.8 | 0.004 | -0.6 | 0.04 | 0.9 | 0.004 |
| **Acetogens** | -0.7 | 0.02 | -0.5 | 0.06 | 0.9 | 0.0005 |

**Table S23.** Spearman correlation coefficients (*ρ*) and corresponding probability (*P*) values determined pair-wise between the relative abundances of different metabolic-types, in the three cores SSK42/5, SSK42/6 and SSK42/9.

|  | **Methanogens** | **Sulfate-reducers** | **ANME** | **Acetogens** | **ANSOB** |  |
| --- | --- | --- | --- | --- | --- | --- |
| **SSK42/5** | | | | | | |
| Methanogens |  | 2.99E-3 | 1.28E-7 | 0.0153 | 0.1825 | ***P* values** |
| Sulfate-reducers | 0.7762 |  | 1.36E-3 | 0.0625 | 0.1825 |  |
| ANME | 0.9720 | 0.8111 |  | 0.0358 | 0.2551 |  |
| Acetogens | 0.6783 | 0.5524 | 0.6083 |  | 0.2453 |  |
| ANSOB | -0.4125 | -0.4126 | -0.3566 | -0.3636 |  |  |
|  | ***ρ* values** | | | |  |  |
| **SSK42/6** | | | | | | |
| Methanogens |  | 5.90E-5 | 1.61E-9 | 1.63E-7 | 0.2466 | ***P* values** |
| Sulfate-reducers | 0.8846 |  | 4.56E-5 | 2.66E-4 | 0.8305 |  |
| ANME | 0.9835 | 0.8901 |  | 2.61E-8 | 0.2233 |  |
| Acetogens | 0.9615 | 0.8461 | 0.9725 |  | 0.1617 |  |
| ANSOB | -0.3461 | -0.0659 | -0.3626 | -0.4120 |  |  |
|  | ***ρ* values** | | | |  |  |
| **SSK42/9** | | | | | | |
| Methanogens |  | 0.1275 | 1.17E-3 | 2.22E-3 | 0.2442 | ***P* values** |
| Sulfate-reducers | 0.5151 |  | 0.1615 | 0.0425 | 0.1869 |  |
| ANME | 0.8667 | 0.4787 |  | 1.63E-3 | 0.4250 |  |
| Acetogens | 0.8424 | 0.6484 | 0.8545 |  | 0.5796 |  |
| ANSOB | -0.4060 | 0.4545 | -0.2848 | -0.2 |  |  |
|  | ***ρ* values** | | | |  |  |

**Table S24.** Spearman correlation coefficients (*ρ*) and corresponding probability (*P*) values calculated pair-wise between the relative abundances of individual sulfur-metabolizing types, concentrations of different sulfur species and sediment-depth, in the three cores SSK42/5, SSK42/6 and SSK42/9.

|  | **Sediment-depth** | **Sulfate concentration** | **Sulfide concentration** | **Prevalence of sulfate-reducers*** | **Prevalence of anaerobic sulfur-oxidizers*** |  |
| --- | --- | --- | --- | --- | --- | --- |
| **SSK42/5** | | | | | |  |
| Sediment-depth |  | 6.1834e-06 | 0.2426 | 0.04 | 0.0348 | ***P* values** |
| Sulfate concentration | -0.9940 |  | 0.2549 | 0.1096 | 0.0488 |  |
| Sulfide concentration | -0.2739 | 0.2671 |  | 0.6092 | 0.1369 |  |
| Prevalence of sulfate-reducers* | -0.6 | 0.4895 | 0.1646 |  | 0.1825 |  |
| Prevalence of anaerobic sulfur-oxidizers* | 0.6224 | -0.5874 | 0.4553 | -0.4126 |  |  |
|  | ***ρ* values** | | | |  |  |
| **SSK42/6** | | | | | |  |
| Sediment-depth |  | 4.0551e-06 | 3.5630e-06 | 0.05 | 0.0655 | ***P* values** |
| Sulfate concentration | -0.9974 |  | 3.4463e-06 | 0.0812 | 0.0655 |  |
| Sulfide concentration | 0.8282 | -0.8288 |  | 0.0183 | 0.0161 |  |
| Prevalence of sulfate-reducers* | -0.5 | 0.5055 | -0.6538 |  | 0.8305 |  |
| Prevalence of anaerobic sulfur-oxidizers* | 0.5818 | -0.5818 | 0.7016 | -0.0659 |  |  |
|  | ***ρ* values** | | | |  |  |
| **SSK42/9** | | | | | |  |
| Sediment-depth |  | 6.0314e-06 | 0.2341 | 0.2 | 0.4697 | ***P* values** |
| Sulfate concentration | -0.9985 |  | 0.2367 | 0.1909 | 0.4697 |  |
| Sulfide concentration | 0.2782 | -0.2767 |  | 0.9186 | 0.2790 |  |
| Prevalence of sulfate-reducers* | 0.5 | -0.4545 | -0.0424 |  | 0.1869 |  |
| Prevalence of anaerobic sulfur-oxidizers* | -0.2606 | 0.2606 | -0.3818 | 0.4545 |  |  |
|  | ***ρ* values** | | | |  |  |

**Table S29.** Sulfate-reducing genera that were found to be most abundant along SSK42/5, according to the taxonomic classification of metagenomic reads (only those entities which had ≥ 5% relative abundance among all sulfate-reducing genera identified in a given sediment-sample have been included).

| **Sediment-depths explored**  **(in cmbsf)** | **Genera identified (Phylum/Class*)** | **Relative abundance among all sulfate-reducing genera detected in the sample (in %)** | **Sulfur compounds reduction phenotype(s) known for the genus** | **Reference(s)** |
| --- | --- | --- | --- | --- |
| 0 | *Desulfovibrio* (Delta) | 14 | Species can reduce SO_4_, S_2_O_3_ as well as S^0^ | (Motamedi and Pedersen 1998) |
|  | *Desulfatibacillum* (Delta) | 12 | Species can reduce SO_4_, SO_3_ as well as S_2_O_3_ | (Cravo-Laureau et al. 2004) |
|  | *Desulfococcus* (Delta) | 9 | Species can reduce SO_4_ and SO_3_ | (Platen et al. 1990) |
|  | *Desulfotomaculum* (Fmt) | 9 | Some species reduce S_2_O_3_, SO_3_ and S^0^ while others reduce SO_4_ | (Hagenauer et al. 1997; Love et al. 1993) |
|  | *Archaeoglobus* (EuA) | 8 | Species can reduce SO_4_, SO_3_ as well as S_2_O_3_ but S^0^ inhibit growth | (Burggraf et al. 1990) |
|  | *Desulfobacterium* (Delta) | 7 | Species can reduce SO_4_, SO_3_ as well as S_2_O_3_ | (Brysch et al. 1987) |
|  | *Desulfitobacterium* (Fmt) | 6 | Species can reduce S^0^, SO_3_ as well as S_2_O_3_ | (Utkin et al. 1994) |
|  | *Desulfuromonas* (Delta) | 5 | Species can reduce S^0^ | (An and Picardal 2015) |
| 15 | *Desulfovibrio* (Delta) | 15 | Species can reduce SO_4_, S_2_O_3_ as well as S^0^ | (Motamedi and Pedersen 1998) |
|  | *Desulfatibacillum* (Delta) | 12 | Species can reduce SO_4_, SO_3_ as well as S_2_O_3_ | (Cravo-Laureau et al. 2004) |
|  | *Desulfococcus* (Delta) | 9 | Species can reduce SO_4_ and SO_3_ | (Platen et al. 1990) |
|  | *Desulfotomaculum* (Fmt) | 9 | Some species reduce S_2_O_3_, SO_3_ and S^0^ while others reduce SO_4_ | (Hagenauer et al. 1997; Love et al. 1993) |
|  | *Archaeoglobus* (EuA) | 9 | Species can reduce SO_4_, SO_3_ as well as S_2_O_3_ but S^0^ inhibit growth | (Burggraf et al. 1990) |
|  | *Desulfobacterium* (Delta) | 7 | Species can reduce SO_4_, SO_3_ as well as S_2_O_3_ | (Brysch et al. 1987) |
|  | *Desulfitobacterium* (Fmt) | 5 | Species can reduce S^0^, SO_3_ as well as S_2_O_3_ | (Utkin et al. 1994) |
|  | *Desulfuromonas* (Delta) | 5 | Species can reduce S^0^ | (An and Picardal 2015) |
| 45 | *Desulfovibrio* (Delta) | 16 | Species can reduce SO_4_, S_2_O_3_ as well as S^0^ | (Motamedi and Pedersen 1998) |
|  | *Desulfatibacillum* (Delta) | 10 | Species can reduce SO_4_, SO_3_ as well as S_2_O_3_ | (Cravo-Laureau et al. 2004) |
|  | *Desulfotomaculum* (Fmt) | 9 | Some species reduce S_2_O_3_, SO_3_ and S^0^ while others reduce SO_4_ | (Hagenauer et al. 1997; Love et al. 1993) |
|  | *Archaeoglobus* (EuA) | 7 | Species can reduce SO_4_, SO_3_ as well as S_2_O_3_ but S^0^ inhibit growth | (Burggraf et al. 1990) |
|  | *Desulfococcus* (Delta) | 6 | Species can reduce SO_4_ and SO_3_ | (Platen et al. 1990) |
|  | *Desulfobacterium* (Delta) | 6 | Species can reduce SO_4_, SO_3_ as well as S_2_O_3_ | (Brysch et al. 1987) |
|  | *Desulfitobacterium* (Fmt) | 6 | Species can reduce S^0^, SO_3_ as well as S_2_O_3_ | (Utkin et al. 1994) |
|  | *Desulfuromonas* (Delta) | 6 | Species can reduce S^0^ | (An and Picardal 2015) |
|  | *Desulfotalea* (Delta) | 6 | Species can reduce SO_4_, SO_3_ as well as S_2_O_3_ | (Knoblauch et al. 1999) |
|  | *Desulfurivibrio* (Delta) | 6 | Species can reduce S^0^, polysulfides & S_2_O_3_ but not sulfate | (Sorokin et al. 2008) |
| 60 | *Desulfovibrio* (Delta) | 19 | Species can reduce SO_4_, S_2_O_3_ as well as S^0^ | (Motamedi and Pedersen 1998) |
|  | *Desulfatibacillum* (Delta) | 10 | Species can reduce SO_4_, SO_3_ as well as S_2_O_3_ | (Cravo-Laureau et al. 2004) |
|  | *Desulfotomaculum* (Fmt) | 8 | Some species reduce S_2_O_3_, SO_3_ and S^0^ while others reduce SO_4_ | (Hagenauer et al. 1997; Love et al. 1993) |
|  | *Desulfococcus* (Delta) | 7 | Species can reduce SO_4_ and SO_3_ | (Platen et al. 1990) |
|  | *Desulfobacterium* (Delta) | 6 | Species can reduce SO_4_, SO_3_ as well as S_2_O_3_ | (Brysch et al. 1987) |
|  | *Archaeoglobus* (EuA) | 6 | Species can reduce SO_4_, SO_3_ as well as S_2_O_3_ but S^0^ inhibit growth | (Burggraf et al. 1990) |
|  | *Desulfuromonas* (Delta) | 6 | Species can reduce S^0^ | (An and Picardal 2015) |
|  | *Desulfotalea* (Delta) | 5 | Species can reduce SO_4_, SO_3_ as well as S_2_O_3_ | (Knoblauch et al. 1999) |
|  | *Desulfitobacterium* (Fmt) | 5 | Species can reduce S^0^, SO_3_ as well as S_2_O_3_ | (Utkin et al. 1994) |
| 90 | *Desulfovibrio* (Delta) | 16 | Species can reduce SO_4_, S_2_O_3_ as well as S^0^ | (Motamedi and Pedersen 1998) |
|  | *Desulfatibacillum* (Delta) | 10 | Species can reduce SO_4_, SO_3_ as well as S_2_O_3_ | (Cravo-Laureau et al. 2004) |
|  | *Desulfuromonas* (Delta) | 9 | Species can reduce S^0^ | (An and Picardal 2015) |
|  | *Desulfotomaculum* (Fmt) | 8 | Some species reduce S_2_O_3_, SO_3_ and S^0^ while others reduce SO_4_ | (Hagenauer et al. 1997; Love et al. 1993) |
|  | *Desulfurivibrio* (Delta) | 7 | Species can reduce S^0^, polysulfides & S_2_O_3_ but not sulfate | (Sorokin et al. 2008) |
|  | *Desulfococcus* (Delta) | 6 | Species can reduce SO_4_ and SO_3_ | (Platen et al. 1990) |
|  | *Desulfobacterium* (Delta) | 6 | Species can reduce SO_4_, SO_3_ as well as S_2_O_3_ | (Brysch et al. 1987) |
|  | *Desulfotalea* (Delta) | 6 | Species can reduce SO_4_, SO_3_ as well as S_2_O_3_ | (Knoblauch et al. 1999) |
|  | *Archaeoglobus* (EuA) | 5 | Species can reduce SO_4_, SO_3_ as well as S_2_O_3_ but S^0^ inhibit growth | (Burggraf et al. 1990) |
| 120 | *Desulfovibrio* (Delta) | 20 | Species can reduce SO_4_, S_2_O_3_ as well as S^0^ | (Motamedi and Pedersen 1998) |
|  | *Desulfatibacillum* (Delta) | 9 | Species can reduce SO_4_, SO_3_ as well as S_2_O_3_ | (Cravo-Laureau et al. 2004) |
|  | *Desulfuromonas* (Delta) | 8 | Species can reduce S^0^ | (An and Picardal 2015) |
|  | *Desulfotomaculum* (Fmt) | 8 | Some species reduce S_2_O_3_, SO_3_ and S^0^ while others reduce SO_4_ | (Hagenauer et al. 1997; Love et al. 1993) |
|  | *Desulfococcus* (Delta) | 7 | Species can reduce SO_4_ and SO_3_ | (Platen et al. 1990) |
|  | *Desulfotalea* (Delta) | 6 | Species can reduce SO_4_, SO_3_ as well as S_2_O_3_ | (Knoblauch et al. 1999) |
|  | *Archaeoglobus* (EuA) | 6 | Species can reduce SO_4_, SO_3_ as well as S_2_O_3_ but S^0^ inhibit growth | (Burggraf et al. 1990) |
|  | *Desulfurivibrio* (Delta) | 6 | Species can reduce S^0^, polysulfides & S_2_O_3_ but not sulfate | (Sorokin et al. 2008) |
|  | *Desulfobacterium* (Delta) | 5 | Species can reduce SO_4_, SO_3_ as well as S_2_O_3_ | (Brysch et al. 1987) |
| 140 | *Desulfurivibrio* (Delta) | 24 | Species can reduce S^0^, polysulfides & S_2_O_3_ but not sulfate | (Sorokin et al. 2008) |
|  | *Desulfotalea* (Delta) | 16 | Species can reduce SO_4_, SO_3_ as well as S_2_O_3_ | (Knoblauch et al. 1999) |
|  | *Desulfobulbus* (Delta) | 10 | Species can reduce SO_4_, SO_3_ as well as S_2_O_3_ | (Suzuki et al. 2007; Sorokin et al. 2012) |
|  | *Desulfovibrio* (Delta) | 9 | Species can reduce SO_4_, S_2_O_3_ as well as S^0^ | (Motamedi and Pedersen 1998) |
|  | *Desulfatibacillum* (Delta) | 6 | Species can reduce SO_4_, SO_3_ as well as S_2_O_3_ | (Cravo-Laureau et al. 2004) |
|  | *Desulfuromonas* (Delta) | 5 | Species can reduce S^0^ | (An and Picardal 2015) |
|  | *Desulfotomaculum* (Fmt) | 5 | Some species reduce S_2_O_3_, SO_3_ and S^0^ while others reduce SO_4_ | (Hagenauer et al. 1997; Love et al. 1993) |
|  | *Desulfobacterium* (Delta) | 5 | Species can reduce SO_4_, SO_3_ as well as S_2_O_3_ | (Brysch et al. 1987) |
| 160 | *Desulfovibrio* (Delta) | 16 | Species can reduce SO_4_, S_2_O_3_ as well as S^0^ | (Motamedi and Pedersen 1998) |
|  | *Desulfuromonas* (Delta) | 13 | Species can reduce S^0^ | (An and Picardal 2015) |
|  | *Desulfotalea* (Delta) | 10 | Species can reduce SO_4_, SO_3_ as well as S_2_O_3_ | (Knoblauch et al. 1999) |
|  | *Desulfurivibrio* (Delta) | 9 | Species can reduce S^0^, polysulfides & S_2_O_3_ but not sulfate | (Sorokin et al. 2008) |
|  | *Desulfatibacillum* (Delta) | 8 | Species can reduce SO_4_, SO_3_ as well as S_2_O_3_ | (Cravo-Laureau et al. 2004) |
|  | *Desulfobacterium* (Delta) | 6 | Species can reduce SO_4_, SO_3_ as well as S_2_O_3_ | (Brysch et al. 1987) |
|  | *Desulfococcus* (Delta) | 6 | Species can reduce SO_4_ and SO_3_ | (Platen et al. 1990) |
|  | *Desulfobulbus* (Delta) | 5 | Species can reduce SO_4_, SO_3_ as well as S_2_O_3_ | (Suzuki et al. 2007; Sorokin et al. 2012) |
|  | *Desulfotomaculum* (Fmt) | 5 | Some species reduce S_2_O_3_, SO_3_ and S^0^ while others reduce SO_4_ | (Hagenauer et al. 1997; Love et al. 1993) |
| 190 | *Desulfovibrio* (Delta) | 18 | Species can reduce SO_4_, S_2_O_3_ as well as S^0^ | (Motamedi and Pedersen 1998) |
|  | *Desulfuromonas* (Delta) | 13 | Species can reduce S^0^ | (An and Picardal 2015) |
|  | *Desulfatibacillum* (Delta) | 9 | Species can reduce SO_4_, SO_3_ as well as S_2_O_3_ | (Cravo-Laureau et al. 2004) |
|  | *Desulfotomaculum* (Fmt) | 9 | Some species reduce S_2_O_3_, SO_3_ and S^0^ while others reduce SO_4_ | (Hagenauer et al. 1997; Love et al. 1993) |
|  | *Archaeoglobus* (EuA) | 6 | Species can reduce SO_4_, SO_3_ as well as S_2_O_3_ but S^0^ inhibit growth | (Burggraf et al. 1990) |
|  | *Desulfitobacterium* (Fmt) | 6 | Species can reduce S^0^, SO_3_ as well as S_2_O_3_ | (Utkin et al. 1994) |
|  | *Desulfobacterium* (Delta) | 6 | Species can reduce SO_4_, SO_3_ as well as S_2_O_3_ | (Brysch et al. 1987) |
|  | *Desulfotalea* (Delta) | 6 | Species can reduce SO_4_, SO_3_ as well as S_2_O_3_ | (Knoblauch et al. 1999) |
|  | *Desulfococcus* (Delta) | 5 | Species can reduce SO_4_ and SO_3_ | (Platen et al. 1990) |
| 220 | *Desulfovibrio* (Delta) | 16 | Species can reduce SO_4_, S_2_O_3_ as well as S^0^ | (Motamedi and Pedersen 1998) |
|  | *Desulfuromonas* (Delta) | 10 | Species can reduce S^0^ | (An and Picardal 2015) |
|  | *Desulfatibacillum* (Delta) | 10 | Species can reduce SO_4_, SO_3_ as well as S_2_O_3_ | (Cravo-Laureau et al. 2004) |
|  | *Desulfobacterium* (Delta) | 9 | Species can reduce SO_4_, SO_3_ as well as S_2_O_3_ | (Brysch et al. 1987) |
|  | *Desulfotomaculum* (Fmt) | 9 | Some species reduce S_2_O_3_, SO_3_ and S^0^ while others reduce SO_4_ | (Hagenauer et al. 1997; Love et al. 1993) |
|  | *Desulfococcus* (Delta) | 6 | Species can reduce SO_4_ and SO_3_ | (Platen et al. 1990) |
|  | *Desulfotalea* (Delta) | 6 | Species can reduce SO_4_, SO_3_ as well as S_2_O_3_ | (Knoblauch et al. 1999) |
|  | *Desulfurivibrio* (Delta) | 6 | Species can reduce S^0^, polysulfides & S_2_O_3_ but not sulfate | (Sorokin et al. 2008) |
| 260 | *Desulfovibrio* (Delta) | 15 | Species can reduce SO_4_, S_2_O_3_ as well as S^0^ | (Motamedi and Pedersen 1998) |
|  | *Desulfurivibrio* (Delta) | 14 | Species can reduce S^0^, polysulfides & S_2_O_3_ but not sulfate | (Sorokin et al. 2008) |
|  | *Desulfotalea* (Delta) | 12 | Species can reduce SO_4_, SO_3_ as well as S_2_O_3_ | (Knoblauch et al. 1999) |
|  | *Desulfuromonas* (Delta) | 10 | Species can reduce S^0^ | (An and Picardal 2015) |
|  | *Desulfatibacillum* (Delta) | 8 | Species can reduce SO_4_, SO_3_ as well as S_2_O_3_ | (Cravo-Laureau et al. 2004) |
|  | *Desulfobulbus* (Delta) | 7 | Species can reduce SO_4_, SO_3_ as well as S_2_O_3_ | (Suzuki et al. 2007; Sorokin et al. 2012) |
|  | *Desulfobacterium* (Delta) | 6 | Species can reduce SO_4_, SO_3_ as well as S_2_O_3_ | (Brysch et al. 1987) |
| 295 | *Desulfotalea* (Delta) | 15 | Species can reduce SO_4_, SO_3_ as well as S_2_O_3_ | (Knoblauch et al. 1999) |
|  | *Desulfurivibrio* (Delta) | 15 | Species can reduce S^0^, polysulfides & S_2_O_3_ but not sulfate | (Sorokin et al. 2008) |
|  | *Desulfovibrio* (Delta) | 13 | Species can reduce SO_4_, S_2_O_3_ as well as S^0^ | (Motamedi and Pedersen 1998) |
|  | *Desulfuromonas* (Delta) | 10 | Species can reduce S^0^ | (An and Picardal 2015) |
|  | *Desulfatibacillum* (Delta) | 7 | Species can reduce SO_4_, SO_3_ as well as S_2_O_3_ | (Cravo-Laureau et al. 2004) |
|  | *Desulfobacterium* (Delta) | 6 | Species can reduce SO_4_, SO_3_ as well as S_2_O_3_ | (Brysch et al. 1987) |

* EuA = Phylum *Euryarchaeota*; Fmt = Phylum *Firmicutes* and Delta = Class *Deltaproteobacteria*.

**Table S30.** Sulfate-reducing genera that were found to be most abundant along SSK42/6, according to the taxonomic classification of metagenomic reads (only those entities which had ≥ 5% relative abundance among all sulfate-reducing genera identified in a given sediment-sample have been included).

| **Sediment-depths explored**  **(in cmbsf)** | **Genera identified (Phylum/Class*)** | **Relative abundance among all sulfate-reducing genera detected in the sample**  **(in %)** | **Sulfur compounds reduction phenotype(s) known for the genus** | **Reference(s)** |
| --- | --- | --- | --- | --- |
| 2 | *Desulfovibrio* (Delta) | 16 | Species can reduce SO_4_, S_2_O_3_ as well as S^0^ | (Motamedi and Pedersen 1998) |
|  | *Desulfatibacillum* (Delta) | 12 | Species can reduce SO_4_, SO_3_ as well as S_2_O_3_ | (Cravo-Laureau et al. 2004) |
|  | *Desulfotomaculum* (Fmt) | 9 | Some species reduce S_2_O_3_, SO_3_ and S^0^ while others reduce SO_4_ | (Hagenauer et al. 1997; Love et al. 1993) |
|  | *Desulfococcus* (Delta) | 8 | Species can reduce SO_4_ and SO_3_ | (Platen et al. 1990) |
|  | *Archaeoglobus* (EuA) | 8 | Species can reduce SO_4_, SO_3_ as well as S_2_O_3_ but S^0^ inhibit growth | (Burggraf et al. 1990) |
|  | *Desulfobacterium* (Delta) | 7 | Species can reduce SO_4_, SO_3_ as well as S_2_O_3_ | (Brysch et al. 1987) |
|  | *Desulfuromonas* (Delta) | 6 | Species can reduce S^0^ | (An and Picardal 2015) |
|  | *Desulfitobacterium* (Fmt) | 5 | Species can reduce S^O^, SO_3_ as well as S_2_O_3_ | (Utkin et al. 1994) |
| 30 | *Desulfovibrio* (Delta) | 16 | Species can reduce SO_4_, S_2_O_3_ as well as S^0^ | (Motamedi and Pedersen 1998) |
|  | *Desulfuromonas* (Delta) | 11 | Species can reduce S^0^ | (An and Picardal 2015) |
|  | *Desulfatibacillum* (Delta) | 8 | Species can reduce SO_4_, SO_3_ as well as S_2_O_3_ | (Cravo-Laureau et al. 2004) |
|  | *Desulfotomaculum* (Fmt) | 8 | Some species reduce S_2_O_3_, SO_3_ and S^0^ while others reduce SO_4_ | (Hagenauer et al. 1997; Love et al. 1993) |
|  | *Archaeoglobus* (EuA) | 7 | Species can reduce SO_4_, SO_3_ as well as S_2_O_3_ but S^0^ inhibit growth | (Burggraf et al. 1990) |
|  | *Desulfobacterium* (Delta) | 6 | Species can reduce SO_4_, SO_3_ as well as S_2_O_3_ | (Brysch et al. 1987) |
|  | *Desulfotalea* (Delta) | 5 | Species can reduce SO_4_, SO_3_ as well as S_2_O_3_ | (Knoblauch et al. 1999) |
|  | *Desulfococcus* (Delta) | 5 | Species can reduce SO_4_ and SO_3_ | (Platen et al. 1990) |
|  | *Desulfitobacterium* (Fmt) | 5 | Species can reduce S^0^, SO_3_ as well as S_2_O_3_ | (Utkin et al. 1994) |
| 45 | *Desulfovibrio* (Delta) | 18 | Species can reduce SO_4_, S_2_O_3_ as well as S^0^ | (Motamedi and Pedersen 1998) |
|  | *Desulfuromonas* (Delta) | 15 | Species can reduce S^0^ | (An and Picardal 2015) |
|  | *Desulfotalea* (Delta) | 7 | Species can reduce SO_4_, SO_3_ as well as S_2_O_3_ | (Knoblauch et al. 1999) |
|  | *Desulfatibacillum* (Delta) | 7 | Species can reduce SO_4_, SO_3_ as well as S_2_O_3_ | (Cravo-Laureau et al. 2004) |
|  | *Desulfotomaculum* (Fmt) | 6 | Some species reduce S_2_O_3_, SO_3_ and S^0^ while others reduce SO_4_ | (Hagenauer et al. 1997; Love et al. 1993) |
|  | *Desulfobacterium* (Delta) | 6 | Species can reduce SO_4_, SO_3_ as well as S_2_O_3_ | (Brysch et al. 1987) |
|  | *Desulfococcus* (Delta) | 5 | Species can reduce SO_4_ and SO_3_ | (Platen et al. 1990) |
|  | *Archaeoglobus* (EuA) | 5 | Species can reduce SO_4_, SO_3_ as well as S_2_O_3_ but S^0^ inhibit growth | (Burggraf et al. 1990) |
|  | *Desulfurivibrio* (Delta) | 5 | Species can reduce S^0^, polysulfides & S_2_O_3_ but not sulfate | (Sorokin et al. 2008) |
| 60 | *Desulfovibrio* (Delta) | 17 | Species can reduce SO_4_, S_2_O_3_ as well as S^0^ | (Motamedi and Pedersen 1998) |
|  | *Desulfuromonas* (Delta) | 11 | Species can reduce S^0^ | (An and Picardal 2015) |
|  | *Desulfatibacillum* (Delta) | 8 | Species can reduce SO_4_, SO_3_ as well as S_2_O_3_ | (Cravo-Laureau et al. 2004) |
|  | *Desulfotomaculum* (Fmt) | 8 | Some species reduce S_2_O_3_, SO_3_ and S^0^ while others reduce SO_4_ | (Hagenauer et al. 1997; Love et al. 1993) |
|  | *Desulfotalea* (Delta) | 7 | Species can reduce SO_4_, SO_3_ as well as S_2_O_3_ | (Knoblauch et al. 1999) |
|  | *Desulfobacterium* (Delta) | 6 | Species can reduce SO_4_, SO_3_ as well as S_2_O_3_ | (Brysch et al. 1987) |
|  | *Archaeoglobus* (EuA) | 6 | Species can reduce SO_4_, SO_3_ as well as S_2_O_3_ but S^0^ inhibit growth | (Burggraf et al. 1990) |
|  | *Desulfitobacterium* (Fmt) | 6 | Species can reduce S^0^, SO_3_ as well as S_2_O_3_ | (Utkin et al. 1994) |
|  | *Desulfococcus* (Delta) | 5 | Species can reduce SO_4_ and SO_3_ | (Platen et al. 1990) |
| 75 | *Desulfovibrio* (Delta) | 18 | Species can reduce SO_4_, S_2_O_3_ as well as S^0^ | (Motamedi and Pedersen 1998) |
|  | *Desulfuromonas* (Delta) | 13 | Species can reduce S^0^ | (An and Picardal 2015) |
|  | *Desulfatibacillum* (Delta) | 9 | Species can reduce SO_4_, SO_3_ as well as S_2_O_3_ | (Cravo-Laureau et al. 2004) |
|  | *Desulfotalea* (Delta) | 8 | Species can reduce SO_4_, SO_3_ as well as S_2_O_3_ | (Knoblauch et al. 1999) |
|  | *Desulfobacterium* (Delta) | 6 | Species can reduce SO_4_, SO_3_ as well as S_2_O_3_ | (Brysch et al. 1987) |
|  | *Desulfotomaculum* (Fmt) | 6 | Some species reduce S_2_O_3_, SO_3_ and S^0^ while others reduce SO_4_ | (Hagenauer et al. 1997; Love et al. 1993) |
|  | *Desulfococcus* (Delta) | 5 | Species can reduce SO_4_ and SO_3_ | (Platen et al. 1990) |
|  | *Desulfurivibrio* (Delta) | 5 | Species can reduce S^0^, polysulfides & S_2_O_3_ but not sulfate | (Sorokin et al. 2008) |
| 90 | *Desulfovibrio* (Delta) | 18 | Species can reduce SO_4_, S_2_O_3_ as well as S^0^ | (Motamedi and Pedersen 1998) |
|  | *Desulfuromonas* (Delta) | 14 | Species can reduce S^0^ | (An and Picardal 2015) |
|  | *Desulfatibacillum* (Delta) | 9 | Species can reduce SO_4_, SO_3_ as well as S_2_O_3_ | (Cravo-Laureau et al. 2004) |
|  | *Desulfotalea* (Delta) | 8 | Species can reduce SO_4_, SO_3_ as well as S_2_O_3_ | (Knoblauch et al. 1999) |
|  | *Desulfobacterium* (Delta) | 7 | Species can reduce SO_4_, SO_3_ as well as S_2_O_3_ | (Brysch et al. 1987) |
|  | *Desulfococcus* (Delta) | 5 | Species can reduce SO_4_ and SO_3_ | (Platen et al. 1990) |
|  | *Desulfotomaculum* (Fmt) | 5 | Some species reduce S_2_O_3_, SO_3_ and S^0^ while others reduce SO_4_ | (Hagenauer et al. 1997; Love et al. 1993) |
|  | *Desulfurivibrio* (Delta) | 5 | Species can reduce S^0^, polysulfides & S_2_O_3_ but not sulfate | (Sorokin et al. 2008) |
| 120 | *Desulfovibrio* (Delta) | 17 | Species can reduce SO_4_, S_2_O_3_ as well as S^0^ | (Motamedi and Pedersen 1998) |
|  | *Desulfuromonas* (Delta) | 13 | Species can reduce S^0^ | (An and Picardal 2015) |
|  | *Desulfotalea* (Delta) | 8 | Species can reduce SO_4_, SO_3_ as well as S_2_O_3_ | (Knoblauch et al. 1999) |
|  | *Desulfatibacillum* (Delta) | 8 | Species can reduce SO_4_, SO_3_ as well as S_2_O_3_ | (Cravo-Laureau et al. 2004) |
|  | *Desulfobacterium* (Delta) | 7 | Species can reduce SO_4_, SO_3_ as well as S_2_O_3_ | (Brysch et al. 1987) |
|  | *Desulfotomaculum* (Fmt) | 6 | Some species reduce S_2_O_3_, SO_3_ and S^0^ while others reduce SO_4_ | (Hagenauer et al. 1997; Love et al. 1993) |
|  | *Desulfurivibrio* (Delta) | 6 | Species can reduce S^0^, polysulfides & S_2_O_3_ but not sulfate | (Sorokin et al. 2008) |
|  | *Desulfococcus* (Delta) | 5 | Species can reduce SO_4_ and SO_3_ | (Platen et al. 1990) |
|  | *Archaeoglobus* (EuA) | 5 | Species can reduce SO_4_, SO_3_ as well as S_2_O_3_ but S^0^ inhibit growth | (Burggraf et al. 1990) |
| 135 | *Desulfovibrio* (Delta) | 19 | Species can reduce SO_4_, S_2_O_3_ as well as S^0^ | (Motamedi and Pedersen 1998) |
|  | *Desulfuromonas* (Delta) | 14 | Species can reduce S^0^ | (An and Picardal 2015) |
|  | *Desulfatibacillum* (Delta) | 9 | Species can reduce SO_4_, SO_3_ as well as S_2_O_3_ | (Cravo-Laureau et al. 2004) |
|  | *Desulfobacterium* (Delta) | 7 | Species can reduce SO_4_, SO_3_ as well as S_2_O_3_ | (Brysch et al. 1987) |
|  | *Desulfotalea* (Delta) | 7 | Species can reduce SO_4_, SO_3_ as well as S_2_O_3_ | (Knoblauch et al. 1999) |
|  | *Desulfotomaculum* (Fmt) | 6 | Some species reduce S_2_O_3_, SO_3_ and S^0^ while others reduce SO_4_ | (Hagenauer et al. 1997; Love et al. 1993) |
|  | *Desulfococcus* (Delta) | 5 | Species can reduce SO_4_ and SO_3_ | (Platen et al. 1990) |
|  | *Desulfomicrobium* (Delta) | 5 | Species can reduce SO_4_, SO_3_ & S_2_O_3_ but not S^0^ | (Thevenieau et al. 2007) |
|  | *Desulfurivibrio* (Delta) | 5 | Species can reduce S^0^, polysulfides & S_2_O_3_ but not sulfate | (Sorokin et al. 2008) |
| 175 | *Desulfovibrio* (Delta) | 19 | Species can reduce SO_4_, S_2_O_3_ as well as S^0^ | (Motamedi and Pedersen 1998) |
|  | *Desulfuromonas* (Delta) | 18 | Species can reduce S^0^ | (An and Picardal 2015) |
|  | *Desulfatibacillum* (Delta) | 8 | Species can reduce SO_4_, SO_3_ as well as S_2_O_3_ | (Cravo-Laureau et al. 2004) |
|  | *Desulfobacterium* (Delta) | 7 | Species can reduce SO_4_, SO_3_ as well as S_2_O_3_ | (Brysch et al. 1987) |
|  | *Desulfotalea* (Delta) | 7 | Species can reduce SO_4_, SO_3_ as well as S_2_O_3_ | (Knoblauch et al. 1999) |
|  | *Desulfurivibrio* (Delta) | 7 | Species can reduce S^0^, polysulfides & S_2_O_3_ but not sulfate | (Sorokin et al. 2008) |
|  | *Desulfomicrobium* (Delta) | 5 | Species can reduce SO_4_, SO_3_ & S_2_O_3_ but not S^0^ | (Thevenieau et al. 2007) |
| 220 | *Desulfuromonas* (Delta) | 20 | Species can reduce S^0^ | (An and Picardal 2015) |
|  | *Desulfovibrio* (Delta) | 14 | Species can reduce SO_4_, S_2_O_3_ as well as S^0^ | (Motamedi and Pedersen 1998) |
|  | *Desulfurivibrio* (Delta) | 10 | Species can reduce S^0^, polysulfides & S_2_O_3_ but not sulfate | (Sorokin et al. 2008) |
|  | *Desulfobacterium* (Delta) | 9 | Species can reduce SO_4_, SO_3_ as well as S_2_O_3_ | (Brysch et al. 1987) |
|  | *Desulfatibacillum* (Delta) | 8 | Species can reduce SO_4_, SO_3_ as well as S_2_O_3_ | (Cravo-Laureau et al. 2004) |
|  | *Desulfurispirillum* (Chrys) | 6 | Species can reduce S^0^ & polysulfides | (Sorokin et al. 2007) |
|  | *Desulfotalea* (Delta) | 6 | Species can reduce SO_4_, SO_3_ as well as S_2_O_3_ | (Knoblauch et al. 1999) |
|  | *Desulfomicrobium* (Delta) | 5 | Species can reduce SO_4_, SO_3_ & S_2_O_3_ but not S^0^ | (Thevenieau et al. 2007) |
| 250 | *Desulfovibrio* (Delta) | 18 | Species can reduce SO_4_, S_2_O_3_ as well as S^0^ | (Motamedi and Pedersen 1998) |
|  | *Desulfuromonas* (Delta) | 12 | Species can reduce S^0^ | (An and Picardal 2015) |
|  | *Desulfotomaculum* (Fmt) | 7 | Some species reduce S_2_O_3_, SO_3_ and S^0^ while others reduce SO_4_ | (Hagenauer et al. 1997; Love et al. 1993) |
|  | *Desulfococcus* (Delta) | 6 | Species can reduce SO_4_ and SO_3_ | (Platen et al. 1990) |
|  | *Desulfatibacillum* (Delta) | 6 | Species can reduce SO_4_, SO_3_ as well as S_2_O_3_ | (Cravo-Laureau et al. 2004) |
|  | *Archaeoglobus* (EuA) | 6 | Species can reduce SO_4_, SO_3_ as well as S_2_O_3_ but S^0^ inhibit growth | (Burggraf et al. 1990) |
|  | *Desulfobacterium* (Delta) | 5 | Species can reduce SO_4_, SO_3_ as well as S_2_O_3_ | (Brysch et al. 1987) |
|  | *Desulfohalobium* (Delta) | 5 | All species can reduce SO_4_ & S_2_O_3_ ; some can reduce SO_3_ & S^0^ | (Ollivier et al. 1991) |
|  | *Desulfurispirillum* (Chrys) | 5 | Species can reduce S^0^ & polysulfides | (Sorokin et al. 2007) |
|  | *Desulfotalea* (Delta) | 5 | Species can reduce SO_4_, SO_3_ as well as S_2_O_3_ | (Knoblauch et al. 1999) |
| 265 | *Desulfovibrio* (Delta) | 14 | Species can reduce SO_4_, S_2_O_3_ as well as S^0^ | (Motamedi and Pedersen 1998) |
|  | *Desulfotomaculum* (Fmt) | 11 | Some species reduce S_2_O_3_, SO_3_ and S^0^ while others reduce SO_4_ | (Hagenauer et al. 1997; Love et al. 1993) |
|  | *Archaeoglobus* (EuA) | 11 | Species can reduce SO_4_, SO_3_ as well as S_2_O_3_ but S^0^ inhibit growth | (Burggraf et al. 1990) |
|  | *Desulfatibacillum* (Delta) | 9 | Species can reduce SO_4_, SO_3_ as well as S_2_O_3_ | (Cravo-Laureau et al. 2004) |
|  | *Desulfitobacterium* (Fmt) | 6 | Species can reduce S^0^, SO_3_ as well as S_2_O_3_ | (Utkin et al. 1994) |
|  | *Desulfococcus* (Delta) | 6 | Species can reduce SO_4_ and SO_3_ | (Platen et al. 1990) |
|  | *Desulfobacterium* (Delta) | 6 | Species can reduce SO_4_, SO_3_ as well as S_2_O_3_ | (Brysch et al. 1987) |
| 275 | *Desulfovibrio* (Delta) | 23 | Species can reduce SO_4_, S_2_O_3_ as well as S^0^ | (Motamedi and Pedersen 1998) |
|  | *Desulfuromonas* (Delta) | 13 | Species can reduce S^0^ | (An and Picardal 2015) |
|  | *Desulfobacterium* (Delta) | 6 | Species can reduce SO_4_, SO_3_ as well as S_2_O_3_ | (Brysch et al. 1987) |
|  | *Desulfurivibrio* (Delta) | 6 | Species can reduce S^0^, polysulfides & S_2_O_3_ but not sulfate | (Sorokin et al. 2008) |
|  | *Desulfatibacillum* (Delta) | 6 | Species can reduce SO_4_, SO_3_ as well as S_2_O_3_ | (Cravo-Laureau et al. 2004) |
|  | *Desulfococcus* (Delta) | 5 | Species can reduce SO_4_ and SO_3_ | (Platen et al. 1990) |
|  | *Desulfohalobium* (Delta) | 5 | All species can reduce SO_4_ & S_2_O_3_ ; some can reduce SO_3_ & S^0^ | (Ollivier et al. 1991) |
|  | *Desulfomicrobium* (Delta) | 5 | Species can reduce SO_4_, SO_3_ & S_2_O_3_ but not S^0^ | (Thevenieau et al. 2007) |
|  | *Desulfotalea* (Delta) | 5 | Species can reduce SO_4_, SO_3_ as well as S_2_O_3_ | (Knoblauch et al. 1999) |
|  | *Desulfotomaculum* (Fmt) | 5 | Some species reduce S_2_O_3_, SO_3_ and S^0^ while others reduce SO_4_ | (Hagenauer et al. 1997; Love et al. 1993) |
|  | *Desulfurispirillum* (Chrys) | 5 | Species can reduce S^0^ & polysulfides | (Sorokin et al. 2007) |

* EuA = Phylum *Euryarchaeota*; Fmt = Phylum *Firmicutes*; Delta = Class *Deltaproteobacteria* and Chrys = Phylum *Chrysiogenetes*.

**Table S31.** Methanogenic genera that were found to be most abundant along SSK42/5, according to the taxonomic classification of metagenomic reads (only those entities which had ≥ 5% relative abundance among all methane-producing genera identified in a given sediment-sample have been included).

|  |  |  | **Methanogenic pathway that is present in all members of the genus (indicated by green shade)** | | | |  |
| --- | --- | --- | --- | --- | --- | --- | --- |
| **Sediment-depths explored**  **(in cmbsf)** | **Genus** | **Relative abundance among all methanogenic genera detected in the sample (in %)** | **Hydrogenotrophic** | **Methylotrophic** | **Acetoclastic** | **Formate-utilizing** | **Reference(s)** |
| 0 | *Methanosarcina* | 16 |  |  |  |  | (Oren 2014a) |
|  | *Methanocaldococcus* | 14 |  |  |  |  | (Oren 2014b) |
|  | *Methanothermobacter* | 9 |  |  |  |  | (Oren 2014c) |
|  | *Methanococcus* | 8 |  |  |  |  | (Oren 2014d) |
|  | *Methanococcoides* | 5 |  |  |  |  | (Oren 2014a) |
| 15 | *Methanosarcina* | 15 |  |  |  |  | (Oren 2014a) |
|  | *Methanocaldococcus* | 14 |  |  |  |  | (Oren 2014b) |
|  | *Methanothermobacter* | 10 |  |  |  |  | (Oren 2014c) |
|  | *Methanococcus* | 9 |  |  |  |  | (Oren 2014d) |
|  | *Methanococcoides* | 5 |  |  |  |  | (Oren 2014a) |
| 45 | *Methanosarcina* | 16 |  |  |  |  | (Oren 2014a) |
|  | *Methanocaldococcus* | 13 |  |  |  |  | (Oren 2014b) |
|  | *Methanococcoides* | 10 |  |  |  |  | (Oren 2014a) |
|  | *Methanothermobacter* | 9 |  |  |  |  | (Oren 2014c) |
|  | *Methanococcus* | 7 |  |  |  |  | (Oren 2014d) |
| 60 | *Methanosarcina* | 18 |  |  |  |  | (Oren 2014a) |
|  | *Methanocaldococcus* | 13 |  |  |  |  | (Oren 2014b) |
|  | *Methanothermobacter* | 10 |  |  |  |  | (Oren 2014c) |
|  | *Methanococcus* | 8 |  |  |  |  | (Oren 2014d) |
|  | *Methanococcoides* | 6 |  |  |  |  | (Oren 2014a) |
| 90 | *Methanosarcina* | 18 |  |  |  |  | (Oren 2014a) |
|  | *Methanocaldococcus* | 13 |  |  |  |  | (Oren 2014b) |
|  | *Methanothermobacter* | 10 |  |  |  |  | (Oren 2014c) |
|  | *Methanococcus* | 7 |  |  |  |  | (Oren 2014d) |
|  | *Methanospirillum* | 6 |  |  |  |  | (Oren 2014e) |
| 120 | *Methanosarcina* | 18 |  |  |  |  | (Oren 2014a) |
|  | *Methanocaldococcus* | 14 |  |  |  |  | (Oren 2014b) |
|  | *Methanothermobacter* | 10 |  |  |  |  | (Oren 2014c) |
|  | *Methanococcus* | 7 |  |  |  |  | (Oren 2014d) |
|  | *Methanosaeta* | 6 |  |  |  |  | (Patel and Sprott 1990) |
| 140 | *Methanosarcina* | 16 |  |  |  |  | (Oren 2014a) |
|  | *Methanocaldococcus* | 15 |  |  |  |  | (Oren 2014b) |
|  | *Methanothermobacter* | 10 |  |  |  |  | (Oren 2014c) |
|  | *Methanococcus* | 8 |  |  |  |  | (Oren 2014d) |
|  | *Methanopyrus* | 6 |  |  |  |  | (Oren 2014f) |
| 160 | *Methanosarcina* | 17 |  |  |  |  | (Oren 2014a) |
|  | *Methanocaldococcus* | 12 |  |  |  |  | (Oren 2014b) |
|  | *Methanothermobacter* | 9 |  |  |  |  | (Oren 2014c) |
|  | *Methanococcus* | 8 |  |  |  |  | (Oren 2014d) |
|  | *Methanospirillum* | 7 |  |  |  |  | (Oren 2014e) |
| 190 | *Methanosarcina* | 16 |  |  |  |  | (Oren 2014a) |
|  | *Methanocaldococcus* | 14 |  |  |  |  | (Oren 2014b) |
|  | *Methanothermobacter* | 10 |  |  |  |  | (Oren 2014c) |
|  | *Methanococcus* | 8 |  |  |  |  | (Oren 2014d) |
|  | *Methanosaeta* | 6 |  |  |  |  | (Patel and Sprott 1990) |
| 220 | *Methanosarcina* | 16 |  |  |  |  | (Oren 2014a) |
|  | *Methanocaldococcus* | 14 |  |  |  |  | (Oren 2014b) |
|  | *Methanothermobacter* | 10 |  |  |  |  | (Oren 2014c) |
|  | *Methanococcus* | 7 |  |  |  |  | (Oren 2014d) |
|  | *Methanosaeta* | 6 |  |  |  |  | (Patel and Sprott 1990) |
| 260 | *Methanosarcina* | 16 |  |  |  |  | (Oren 2014a) |
|  | *Methanocaldococcus* | 14 |  |  |  |  | (Oren 2014b) |
|  | *Methanothermobacter* | 9 |  |  |  |  | (Oren 2014c) |
|  | *Methanococcus* | 7 |  |  |  |  | (Oren 2014d) |
|  | *Methanococcoides* | 7 |  |  |  |  | (Oren 2014a) |
| 295 | *Methanococcoides* | 18 |  |  |  |  | (Oren 2014a) |
|  | *Methanosarcina* | 17 |  |  |  |  | (Oren 2014a) |
|  | *Methanosaeta* | 10 |  |  |  |  | (Patel and Sprott 1990) |
|  | *Methanocaldococcus* | 9 |  |  |  |  | (Oren 2014b) |

**Table S32.** Methanogenic genera that were found to be most abundant along SSK42/6, according to the taxonomic classification of metagenomic reads (only those entities which had ≥ 5% relative abundance among all methane-producing genera identified in a given sediment-sample have been included).

|  |  |  | **Methanogenic pathway that is present in all members of the genus (indicated by green shade)** | | | |  |
| --- | --- | --- | --- | --- | --- | --- | --- |
| **Sediment-depths explored**  **(in cmbsf)** | **Genus** | **Relative abundance among all methanogenic genera detected in the sample (in %)** | **Hydrogenotrophic** | **Methylotrophic** | **Acetoclastic** | **Formate-utilizing** | **Reference(s)** |
| 2 | *Methanosarcina* | 16 |  |  |  |  | (Oren 2014a) |
|  | *Methanocaldococcus* | 14 |  |  |  |  | (Oren 2014b) |
|  | *Methanothermobacter* | 9 |  |  |  |  | (Oren 2014c) |
|  | *Methanococcus* | 8 |  |  |  |  | (Oren 2014d) |
|  | *Methanococcoides* | 5 |  |  |  |  | (Oren 2014a) |
| 30 | *Methanosarcina* | 16 |  |  |  |  | (Oren 2014a) |
|  | *Methanocaldococcus* | 14 |  |  |  |  | (Oren 2014b) |
|  | *Methanothermobacter* | 10 |  |  |  |  | (Oren 2014c) |
|  | *Methanococcus* | 7 |  |  |  |  | (Oren 2014d) |
|  | *Methanococcoides* | 5 |  |  |  |  | (Oren 2014a) |
| 45 | *Methanosarcina* | 17 |  |  |  |  | (Oren 2014a) |
|  | *Methanocaldococcus* | 13 |  |  |  |  | (Oren 2014b) |
|  | *Methanothermobacter* | 9 |  |  |  |  | (Oren 2014c) |
|  | *Methanococcus* | 8 |  |  |  |  | (Oren 2014d) |
|  | *Methanosaeta* | 6 |  |  |  |  | (Patel and Sprott 1990) |
| 60 | *Methanosarcina* | 17 |  |  |  |  | (Oren 2014a) |
|  | *Methanocaldococcus* | 14 |  |  |  |  | (Oren 2014b) |
|  | *Methanothermobacter* | 10 |  |  |  |  | (Oren 2014c) |
|  | *Methanococcus* | 8 |  |  |  |  | (Oren 2014d) |
|  | *Methanosaeta* | 6 |  |  |  |  | (Patel and Sprott 1990) |
| 75 | *Methanosarcina* | 19 |  |  |  |  | (Oren 2014a) |
|  | *Methanocaldococcus* | 13 |  |  |  |  | (Oren 2014b) |
|  | *Methanothermobacter* | 9 |  |  |  |  | (Oren 2014c) |
|  | *Methanococcus* | 7 |  |  |  |  | (Oren 2014d) |
|  | *Methanosaeta* | 5 |  |  |  |  | (Patel and Sprott 1990) |
| 90 | *Methanosarcina* | 18 |  |  |  |  | (Oren 2014a) |
|  | *Methanocaldococcus* | 13 |  |  |  |  | (Oren 2014b) |
|  | *Methanothermobacter* | 9 |  |  |  |  | (Oren 2014c) |
|  | *Methanococcus* | 7 |  |  |  |  | (Oren 2014d) |
|  | *Methanosaeta* | 6 |  |  |  |  | (Patel and Sprott 1990) |
| 120 | *Methanosarcina* | 17 |  |  |  |  | (Oren 2014a) |
|  | *Methanocaldococcus* | 13 |  |  |  |  | (Oren 2014b) |
|  | *Methanothermobacter* | 9 |  |  |  |  | (Oren 2014c) |
|  | *Methanococcus* | 8 |  |  |  |  | (Oren 2014d) |
|  | *Methanobrevibacter* | 7 |  |  |  |  | (Oren 2014c) |
| 135 | *Methanosarcina* | 21 |  |  |  |  | (Oren 2014a) |
|  | *Methanocaldococcus* | 13 |  |  |  |  | (Oren 2014b) |
|  | *Methanococcus* | 10 |  |  |  |  | (Oren 2014d) |
|  | *Methanothermobacter* | 8 |  |  |  |  | (Oren 2014c) |
| 175 | *Methanosarcina* | 22 |  |  |  |  | (Oren 2014a) |
|  | *Methanocaldococcus* | 9 |  |  |  |  | (Oren 2014b) |
|  | *Methanococcus* | 8 |  |  |  |  | (Oren 2014d) |
|  | *Methanothermobacter* | 7 |  |  |  |  | (Oren 2014c) |
|  | *Methanobrevibacter* | 7 |  |  |  |  | (Oren 2014c) |
| 220 | *Methanosarcina* | 25 |  |  |  |  | (Oren 2014a) |
|  | *Methanocaldococcus* | 10 |  |  |  |  | (Oren 2014b) |
|  | *Methanothermobacter* | 8 |  |  |  |  | (Oren 2014c) |
|  | *Methanococcus* | 7 |  |  |  |  | (Oren 2014d) |
| 250 | *Methanosarcina* | 15 |  |  |  |  | (Oren 2014a) |
|  | *Methanocaldococcus* | 14 |  |  |  |  | (Oren 2014b) |
|  | *Methanothermobacter* | 10 |  |  |  |  | (Oren 2014c) |
|  | *Methanococcus* | 9 |  |  |  |  | (Oren 2014d) |
|  | *Methanosaeta* | 6 |  |  |  |  | (Patel and Sprott 1990) |
| 265 | *Methanosarcina* | 15 |  |  |  |  | (Oren 2014a) |
|  | *Methanocaldococcus* | 14 |  |  |  |  | (Oren 2014b) |
|  | *Methanothermobacter* | 11 |  |  |  |  | (Oren 2014c) |
|  | *Methanococcus* | 8 |  |  |  |  | (Oren 2014d) |
|  | *Methanopyrus* | 6 |  |  |  |  | (Oren 2014f) |
| 275 | *Methanosarcina* | 19 |  |  |  |  | (Oren 2014a) |
|  | *Methanocaldococcus* | 12 |  |  |  |  | (Oren 2014b) |
|  | *Methanothermobacter* | 9 |  |  |  |  | (Oren 2014c) |
|  | *Methanococcus* | 8 |  |  |  |  | (Oren 2014d) |
|  | *Methanococcoides* | 5 |  |  |  |  | (Oren 2014a) |

**References used in the Supplementary Tables**

An TT, Picardal FW. *Desulfuromonas carbonis* sp. nov., an Fe (III)-, S0-and Mn (IV)-reducing bacterium isolated from an active coalbed methane gas well. *Int J Syst Evol Microbiol* 2015;**65:**1686-93.

Brysch K, Schneider C, Fuchs G et al. Lithoautotrophic growth of sulfate-reducing bacteria, and description of *Desulfobacterium autotrophicum* gen. nov., sp. nov. *Arch Microbiol* 1987;**148:**264-74.

Burggraf S, Jannasch HW, Nicolaus B et al. *Archaeoglobus profundus* sp. nov., represents a new species within the sulfate-reducing Archaebacteria. *Syst Appl Microbiol* 1990;**13:**24-28.

Cravo-Laureau C, Matheron R, Cayol JL et al. *Desulfatibacillum aliphaticivorans* gen. nov., sp. nov., an n-alkane- and n-alkene-degrading, sulfate-reducing bacterium. *Int J Syst Evol Microbiol* 2004;**54:**77-83.

Hagenauer A, Hippe H, Rainey FA. *Desulfotomaculum aeronauticum* sp. nov., a sporeforming, thiosulfate-reducing bacterium from corroded aluminium alloy in an aircraft. *Syst Appl Microbiol* 1997;**20:**65-71.

Knoblauch C, Sahm K, Jørgensen BB. Psychrophilic sulfate-reducing bacteria isolated from permanently cold Arctic marine sediments: description of *Desulfofrigus oceanense* gen. nov., sp. nov., *Desulfofrigus fragile* sp. nov., *Desulfofaba gelida* gen. nov., sp. nov., *Desulfotalea psychrophila* gen. nov., sp. nov. and *Desulfotalea arctica* sp. nov. *Int J Syst Evol Microbiol* 1999;**49:**1631-43.

Love CA, Patel BKC, Nichols PD et al. *Desulfotomaculum australicum*, sp. nov., a thermophilic sulfate-reducing bacterium isolated from the Great Artesian Basin of Australia. *Syst Appl Microbiol* 1993;**16:**244-51.

Motamedi M, Pedersen K. *Desulfovibrio aespoeensis* sp. nov., a mesophilic sulfate-reducing bacterium from deep groundwater at Aspo hard rock laboratory, Sweden. *Int J Syst Bacteriol* 1998;**48:**311-5.

Ollivier B, Hatchikian CE, Prensier G et al. *Desulfohalobium retbaense* gen. nov., sp. nov., a halophilic sulfate-reducing bacterium from sediments of a hypersaline lake in Senegal. *Int J Syst Evol Microbiol* 1991;**41:**74-81.

Oren A. The family *Methanosarcinaceae*. In: Rosenberg E, DeLong EF, Lory S, Stackebrandt E, Thompson F (eds.). *The Procaryotes*. Heidelberg: Springer, 2014a, 259-281.

Oren A. The family *Methanocaldococcaceae*. In: Rosenberg E, DeLong EF, Lory S, Stackebrandt E, Thompson F (eds.). *The Procaryotes*. Heidelberg: Springer, 2014b, 201-208.

Oren A. The family *Methanobacteriaceae*. In: Rosenberg E, DeLong EF, Lory S, Stackebrandt E, Thompson F (eds.). *The Procaryotes*. Heidelberg: Springer, 2014c, 165-193.

Oren A. The family *Methanococcaceae*. In: Rosenberg E, DeLong EF, Lory S, Stackebrandt E, Thompson F (eds.). *The Procaryotes*. Heidelberg: Springer, 2014d, 215-224.

Oren A. The family *Methanospirillaceae*. In: Rosenberg E, DeLong EF, Lory S, Stackebrandt E, Thompson F (eds.). *The Procaryotes*. Heidelberg: Springer, 2014e, 283-290.

Oren A. The Family *Methanopyraceae*. In: Rosenberg E, DeLong EF, Lory S, Stackebrandt E, Thompson F (eds.). *The Procaryotes*. Heidelberg: Springer, 2014f, 247-252.

Patel GB, Sprott GD. *Methanosaeta concilii* gen. nov., sp. nov.(“*Methanothrix concilii*”) and *Methanosaeta thermoacetophila* nom. rev., comb. nov. *Int J Syst Evol Microbiol* 1990;**40:**79-82.

Platen H, Temmes A, Schink B. Anaerobic degradation of acetone by *Desulfococcus biacutus* spec. nov. *Arch Microbiol* 1990;**154:**355-61.

Sorokin DY, Foti M, Tindall BJ et al. *Desulfurispirillum alkaliphilum* gen. nov. sp. nov., a novel obligately anaerobic sulfur- and dissimilatory nitrate-reducing bacterium from a full-scale sulfide-removing bioreactor. *Extremophiles* 2007;**11:**363-70.

Sorokin DY, Tourova TP, Mußmann M et al. *Dethiobacter alkaliphilus* gen. nov. sp. nov., and *Desulfurivibrio alkaliphilus* gen. nov. sp. nov.: two novel representatives of reductive sulfur cycle from soda lakes. *Extremophiles* 2008;**12:**431-9.

Sorokin DY, Tourova TP, Panteleeva AN et al. *Desulfonatronobacter acidivorans* gen. nov., sp. nov. and *Desulfobulbus alkaliphilus* sp. nov., haloalkaliphilic heterotrophic sulfate-reducing bacteria from soda lakes. *Int J Syst Evol Microbiol* 2012;**62:**2107-13.

Suzuki D, Ueki A, Amaishi A et al. *Desulfobulbus japonicus* sp. nov., a novel gram-negative propionate-oxidizing, sulfate-reducing bacterium isolated from an estuarine sediment in Japan. *Int J Syst Evol Microbiol* 2007;**57:**849-55.

Thevenieau F, Fardeau M-L, Ollivier B et al. *Desulfomicrobium thermophilum* sp. nov., a novel thermophilic sulphate-reducing bacterium isolated from a terrestrial hot spring in Colombia. *Extremophiles* 2007;**11:**295-303.

Utkin I, Woese C, Wiegel J. Isolation and characterization of *Desulfitobacterium dehalogenans* gen. nov., sp. nov., an anaerobic bacterium which reductively dechlorinates chlorophenolic compounds. *Int J Syst Evol Microbiol* 1994;**44:**612-9.

**Supplementary Notes**

**Supplementary Note 1**

Referred citations for the bacterial and archaeal taxa, summing up the mean relative abundances of which gave the measure of prevalence of sulfate-reducers within a sediment community of SSK42/5, SSK42/6 or SSK42/9 (references are cited after the names of the taxa).

1. *Archaeoglobus, Desulfurobacterium, Desulfacinum*, *Desulfobacca*, *Desulfomonile, Desulforhabdus*, *Desulfovibrio*, *Desulfurella*, *Desulfuromonas*, *Desulfuromusa* and *Thermodesulforhabdus, Desulfitobacterium*, *Desulfosporosinus*, *Desulfotomaculum*, *Thermodesulfovibrio, Dethiosulfovibrio, Thermodesulfobacterium* (Rabus et al. 2006);
2. *Desulfurococcus* (Kublanov et al. 2009);
3. *Desulfurolobus* (Zillig et al. 1986);
4. *Desulfurispira* (Sorokin and Muyzer, 2010);
5. *Desulfurispirillum* (Sorokin et al. 2007);
6. *Desulfobaculum* (Zhao et al. 2012);
7. *Desulfocurvus* (Klouche et al. 2009);
8. *Desulfoglaeba* (Davidova et al. 2006);
9. *Desulfomonas* (Moore et al. 1976);
10. *Desulfosoma* (Baena et al. 2011);
11. *Desulfovirga* (Tanaka et al. 2000);
12. *Desulfitibacter* (Nielsen et al. 2006);
13. *Desulfitispora* (Sorokin and Chernyh, 2017);
14. *Desulfonispora* (Denger et al. 1999);
15. *Desulfurispora* (Kaksonen et al. 2007a);
16. *Desulfovirgula­ (*Kaksonen et al. 2007a);
17. *Dethiobacter* (Sorokin et al. 2008);
18. *Dethiosulfatibacter* (Takii et al. 2007);
19. *Thermodesulfobium­* (Frolov et al. 2017);
20. *Thermodesulfatator* (Lai et al. 2016);
21. *Desulfarculaceae* (Kuever, 2014a);
22. *Desulfobacteraceae* (Kuever, 2014b);
23. *Desulfobulbaceae­* (Kuever, 2014c);
24. *Desulfohalobiaceae* (Kuever, 2014d);
25. *Desulfomicrobiaceae* (Kuever and Galushko, 2014);
26. *Desulfonatronaceae* (Kuever, 2014e).

**Complete References**

**Supplementary Note 2**

Referred citations for the anaerobically sulfur-oxidizing chemolithotrophic bacteria (ANSOB), summing up the mean relative abundances of which gave the measure of prevalence of ANSOB within a sediment community of SSK42/5, SSK42/6 or SSK42/9 (references are cited after the names of the taxa).

1. *Beggiatoa* (Dubinina et al. 2017);
2. *Sulfuricurvum* (Kodama et al. 2004);
3. *Sulfurimonas* (Labrenz et al. 2013);
4. *Sulfurovum* (Mori et al. 2018);
5. *Thiobacillus* (Kellermann et al. 2009).
6. *Thioploca* (Maier and Gallardo, 1984)
7. *Thiomargarita* (Schulz et al. 1999)

**Complete References**
